# Proliferation of Early and Mature Osteoblasts Is Required for Bone Fracture Healing in Mice

**DOI:** 10.1101/2025.05.27.656371

**Authors:** Nicole R. Gould, Andre F. Coello, Jennifer A. McKenzie, Tiandao Li, Katherine R. Hixon, Leyi Chen, Kristen Barwick, Tiffany Lee, Mariam Obaji, Bo Zhang, David M. Ornitz, Matthew J. Silva

## Abstract

Fractures heal by rapid formation of mineralized callus. Essential to this process is proliferation of periosteal cells to supply bone-forming osteoblasts. To better understand the role of cell proliferation in fracture healing, we asked: How is callus composition altered when osteolineage cells proliferation is blocked? Do mature osteoblasts proliferate to contribute to callus formation? First, mice expressing herpes simplex virus-thymidine kinase (HSV-TK) driven by the 3.6Col1a1 promoter were treated with ganciclovir (GCV) to ablate proliferating osteolineage cells for 5 or 10 days. Analysis of callus cells using single-cell RNA-seq revealed that GCV-treated Col1-TK mice had fewer osteoblasts and chondrocytes than control mice, with more myofibroblasts and immune cells, consistent with fibrous nonunion. In controls, 15-30% of callus cells that expressed the early osteoblast markers osterix (OSX) and the late marker osteocalcin (OCN) were in the cell cycle. Next, we targeted proliferating osteoblasts at different stages of differentiation by crossing Osx-CreERT2, Ocn-Cre and Dmp1-CreERT2 mice with novel ROSA-TK mice. Following fracture, each Cre;ROSA-TK mouse line exhibited poorer radiographic healing, decreased callus bone volume and a shift from callus bone to fibrous tissue. We conclude that osteoblasts, often considered post-mitotic, proliferate after fracture to contribute to formation of mineralized callus essential to healing.

## Introduction

Healing of broken bones occurs via a regenerative process involving multiple cell types and overlapping stages of inflammation, fibrovascular callus formation, intramembranous and endochondral ossification, and bone remodeling (1, 2). Functionally, long bone healing is successful when a periosteal mineralized callus bridges the fracture gap with sufficient stiffness and strength to allow weight bearing. Failed healing leads to fracture nonunion, characterized by inadequate osteochondral tissue and persistence of fibrovascular tissue (3, 4). In recent years, our understanding of cell types and molecular factors that contribute to fracture healing has increased greatly with the use of mouse genetic and lineage tracing models. In particular, the contribution to callus formation of different skeletal stem and progenitor cell (SSPC) populations has been described (5–8).

Among the cell types essential for fracture healing is the osteoblast. Bone-forming osteoblasts are needed in great number for mineralized callus formation, and rapid expansion of this population must involve proliferation of osteoprogenitors and/or more differentiated cells. Proliferation of periosteal cells is a histological hallmark of early fracture healing (9–11). However, the molecular identity of these proliferating cells is not well described. In particular, the differentiation stage of osteolineage cells at the time of their proliferation is unknown. Osteolineage cell proliferation in fracture healing is often assumed to occur at the progenitor stage (1, 2, 7), consistent with a notion of the mature osteoblast as post-mitotic cell (7, 12–14). This view is based on classic *in vitro* studies (12, 15) and histological observations of cells in developing animals (16, 17), but has not been critically examined in the context of fracture healing.

We recently studied the role of osteolineage cell proliferation in fracture healing using the Col1-TK mouse, in which herpes simplex virus (HSV) thymidine kinase (TK) expression is driven by the 3.6-kb fragment of the *Col1a1* promoter (18). Treatment with ganciclovir, a nucleoside analog that incorporates into the DNA of replicating cells and is converted to a toxic form by HSV-TK, leads to ablation of proliferating 3.6Col1a1 lineage cells which in turn results in minimal callus formation and fracture nonunion (19). This finding demonstrated that proliferation of osteolineage cells is essential to fracture healing. But because the 3.6Col1a1 promoter is active in both pre- and mature osteoblasts and periosteal fibroblasts (20), it remains unclear at what stage(s) of differentiation are osteolineage cells proliferating to contribute to fracture healing.

Here we characterize the cellular composition of the early fracture callus in normal healing mice and in Col1-TK non-healers, with the goal of identifying the contribution of early and mature osteoblast proliferation to mineralized callus formation. Using single cell mRNA sequencing (scRNAseq) we identify a shift from osteochondral cells to fibroblasts in normal versus non-healing mice. Using a novel Cre-inducible ROSA-TK mouse we show that proliferation of both early and mature osteoblasts is essential to fracture healing.

## Results

### Ablation of proliferating 3.6Col1a1-lineage cells for the first 7 or 14 days after fracture impairs callus bone formation

When Col1-TK mice are administered ganciclovir (GCV), HSV-TK expressed in 3.6Col1a1-lineage cells converts GCV into a toxic form, which incorporates into the DNA of replicating cells to induce apoptosis (18). We reported that GCV dosing for 14 or 28 days following fracture in Col1-TK mice results in failure of mineralized callus formation, establishing the first 2 weeks as a critical time for osteogenic cell proliferation (19). To extend the prior work, we treated WT control and TK+ experimental mice with GCV for 3, 7, or 14 days and evaluated fracture healing at 21 days post-fracture (DPF) (Fig 1A). Based on radiographic analysis, the periosteal fracture callus in 90-100% of control mice was fully bridged by 21 DPF, regardless of GCV dosing duration (Fig 1B, Suppl Fig S1A). Experimental mice treated with GCV for 3 days had delayed callus formation compared to control, but by 21 DPF 100% had a fully bridged callus. In contrast, experimental mice treated with GCV for 7 or 14 days had significantly impaired bridging, with only 57% and 14% of samples fully bridged, respectively.

**Figure 1:**
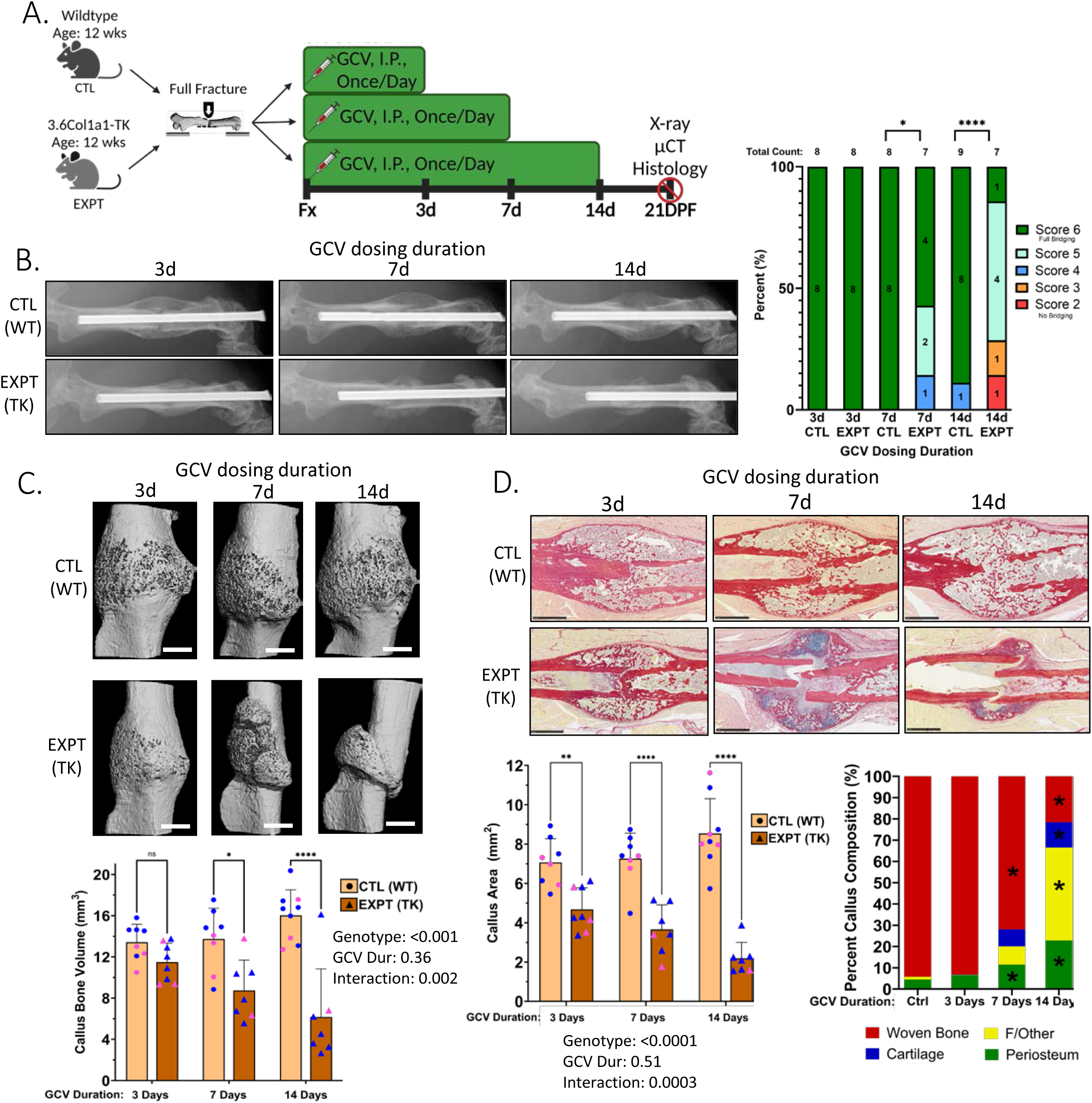
Ablation of proliferating 3.6Col1a1-lineage cells in the first 3-14 days following fracture causes defects in callus formation at 21 days post-fracture. **(A)** Femur fractures were created in 12-week-old wildtype control (CTL) and Col1-TK experimental (EXPT) mice, followed by ganciclovir (GCV) dosing for 3, 7, or 14 days (n=7-9). **(B)** (Left) Representative radiographs at 21 days post-fracture (DPF). (Right) Quantification of radiographic healing. **(C)** 3D microCT images at 21 DPF (scale bar, 1 mm). **(D)** Sagittal callus sections were stained with picrosirius red (PSR) to visualize collagen-rich bone and alcian blue (AB) for proteoglycan-rich cartilage, and callus composition was determined (scale bar=1 mm). (Graphs depict mean ± SD; individual data points shown [pink=female; blue=male]. Statistics: Chi-square test (B) or two-Way ANOVA with Tukey Post Hoc test (C,D); *p<0.05, **<0.01, ****<0.0001; ns: p>0.05)

Consistent with radiographic findings, microCT of fracture callus showed that experimental mice treated with GCV for 7 or 14 days had progressively less bone volume at 21 DPF compared to controls, while 3 days of GCV did not alter callus bone volume (Fig 1C). Other microCT parameters also showed a significant interaction between GCV duration and genotype, with longer treatment durations impairing callus formation in experimental but not control mice (Suppl Fig S1B). Notably, the parameter most affected by 3 days GCV dosing was total volume, indicating that targeting proliferation for the first 3 days of fracture healing reduces eventual callus size at 21 DPF.

Histological analysis also demonstrated progressively smaller callus size and altered tissue composition in experimental mice with longer GCV dosing (Fig 1D, Suppl Fig S1C). At 21 DPF, in both control and experimental mice treated with GCV for 3 days, callus composition was 95% woven bone, indicating that healing had progressed past the cartilage stage. In contrast, GCV treatment for 7 or 14 days in experimental mice significantly altered callus composition, with woven bone comprising 70% and 20% of the callus and fibrous tissue comprising 8% and 45% of the callus, respectively. In summary, ablation of proliferating osteoblast-lineage cells for the first 3 days after fracture resulted in modest, transient delays in healing, while ablation for 7 days led to moderate impairments in callus size and composition at 21 DPF, and ablation for 14 days resulted in a fracture callus with markedly less bone and more fibrous tissue.

### 5-day ablation of proliferating 3.6Col1a1-lineage cells depletes fracture callus of osteoblasts and chondrocytes

As ablation of proliferating 3.6Col1a1-lineage cells for 7 or 14 days after fracture caused persistent impairment in callus size and composition, we sought to profile the underlying changes in the early callus at the single-cell level. Following fracture, Col1-TK mice were treated either with water (Control) or GCV (Experimental) for 5 days (Fig 2A). Periosteal whole-callus tissue was dissected and enzymatically digested, and the resulting cell samples (were profiled using scRNAseq. For analysis, Control and Experimental samples were integrated before clustering to facilitate comparisons between groups. Unbiased cluster analysis revealed that the early fracture callus contains multiple cell types of three major lineages: mesenchymal (*Prrx1*+), endothelial (*Pecam1*+), and immune (*Ptprc*/CD45+) (Fig 2B, Suppl Fig S2A,B).

**Figure 2:**
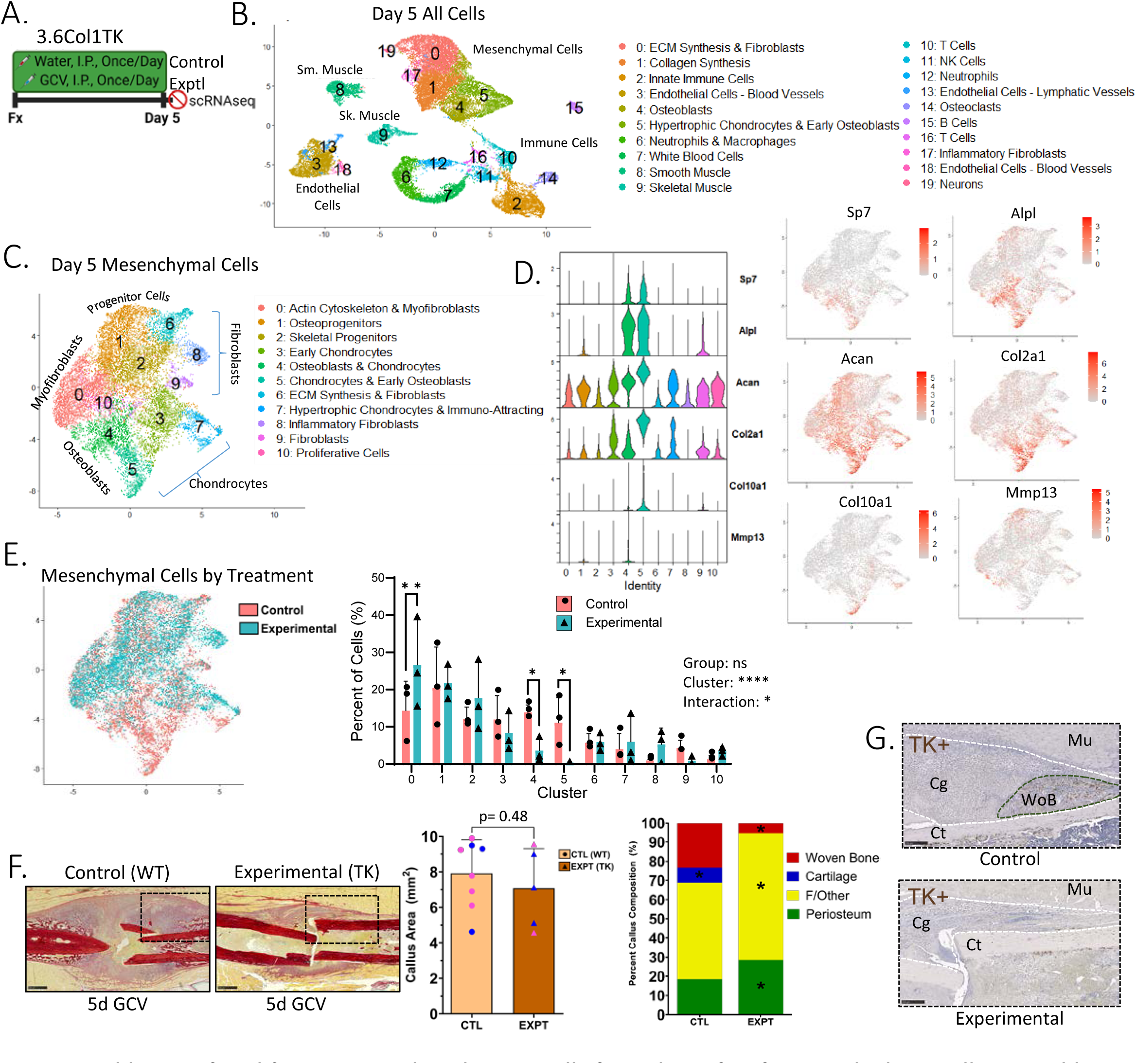
Ablation of proliferating 3.6Col1a1 lineage cells for 5 days after fracture depletes callus osteoblasts and chondrocytes and alters callus composition. **(A)** After fracture Col1-TK mice were treated with water (Control, n=3) or GCV (Exptl, n=3) for 5 days. Cells isolated from whole-fracture callus were subjected to scRNAseq. **(B)** UMAP plot of total callus cells with annotated cell cluster identities based on canonical gene expression and bioinformatic analysis (COMPBIO). **(C)** Mesenchymal cells were selected *in silico* and re-clustered. **(D)** Expression of canonical genes for osteoblasts (*Sp7*, *Alpl*), chondrocytes (*Acan*, *Col2a1*), and hypertrophic chondrocytes (*Col10a1*, *Mmp13*) projected on UMAPS and displayed as violin plots. **(E)** Mesenchymal cells were identified by treatment group, and the percent of cells in each cluster was quantified (n=3). **(F)** Histological sections of callus at 5 DPF from Control (n=8; WT + GCV, or Col1-TK + water) and Exptl (n=5; Col1-TK + GCV) were stained with picrosirius red and alcian blue (PSRAB) and used to determine total callus area, and percent area of woven bone (red), cartilage (blue), fibrous/other tissue, and periosteum. **(G)** High-power fields (from F) of TK IHC of control (Col1-TK + water) and experimental (Col1-TK + GCV) mice (white dashed line=callus periphery, green dashed line=woven bone area, Mu=muscle, Cg=cartilage, Ct=Cortical bone, WoB=woven bone). (Statistics: two-way ANOVA with Holm-Sidak Post Hoc test (E) or Two-Tailed t-test (F). Scale bars: PSRAB=0.5 mm, TK=1 mm.)

Notably, hypertrophic chondrocytes and early osteoblasts were depleted in Experimental mice; these cells (Cluster 5) comprised 12% of all callus cells in Control mice, but only 1% in Experimental samples (Suppl Fig S2C).

To enhance resolution of the mesenchymal population, cells of the original mesenchymal clusters (0, 1, 4, 5, 17) were selected *in silico* for re-clustering. Analysis of this mesenchymal sub-population identified 11 clusters; we assigned cell identities to each cluster based on canonical gene expression and bioinformatic analysis of differentially expressed genes. Clusters were comprised of skeletal progenitors, fibroblasts, chondrocytes, and osteoblasts (Figs 2C, D). Osteoblasts (Cluster 4) and chondrocytes (Cluster 5) were significantly depleted in Experimental mice (4% of all cells) compared to Control mice (25% of all cells) (Fig 2E). On the other hand, cells with high expression of actin cytoskeletal genes that resemble myofibroblasts (Cluster 0) were significantly enriched in the Experimental fracture callus (CTL: 14%, EXPT: 27%). The cells in Cluster 8 expressed non-specific fibroblast markers (*Col1a1*, *Col3a1*, *Postn*) as well as strong and specific expression of markers of an innate immune response (*Isg15*, *Ifit1*, *Irf7*), and were labeled “inflammatory fibroblasts” (Suppl Fig S2D).

We also performed histological analysis at 5 DPF, when the periosteal callus is comprised mainly of fibrous/undifferentiated tissue, with small amounts of woven bone and early cartilage. Callus area did not differ between Control and Experimental groups, but callus composition was altered; callus tissue from Experimental mice had less woven bone and cartilage and more fibrous and periosteal tissue (Fig 2F, Suppl Fig S2E). Immunostaining for HSV-TK confirmed the loss of TK+ cells in calluses of Experimental (GCV-treated Col1-TK) mice compared to Control (water-treated Col1-TK) mice, most notably in areas of woven bone (Fig 2G). Thus, scRNAseq and histology indicate that ablation of proliferating 3.6Col1a1-lineage cells in the first 5 days after fracture deprives the callus of osteoblasts and chondrocytes, resulting in less cartilage and bone formation and a shift toward less differentiated fibrous tissue.

### 10-day ablation of proliferating 3.6Col1a1-lineage cells depletes fracture callus of osteoblasts and chondrocytes, and causes a shift toward fibroblasts and fibrous tissue

We next analyzed the callus at 10 DPF, when osteochondral cells and tissue are normally predominant (Fig 3A). scRNAseq of total callus cells again revealed a mixed population of mesenchymal, endothelial, and immune lineages (Fig 3B, Suppl Fig S3A,B). The proportion of cells in the mesenchymal clusters at 10 DPF (69% of all cells) was greater than at 5 DPF (46%), consistent with callus maturation. In calluses from Control mice, 33% of all cells were in Clusters 1 (mature chondrocytes) and 2 (hypertrophic chondrocytes & osteoblasts). In contrast, in calluses from Experimental mice, only 9% of cells were in these two clusters, indicating significant depletion of mature osteochondral cells (Suppl Fig S3C). The opposite shift was seen in cells in Clusters 5 (smooth muscle cells) and 6 (inflammatory fibroblasts), which were enriched in Experimental mice (21% of all cells) compared to Control (4.4%). In Control mice, the number of cells in immune clusters was reduced by 71% from 5 to 10 DPF (p=0.04, Fig 3C), indicating significant resolution of the early immune cell response. In contrast, the total number of cells in immune clusters was reduced by only 43% during this interval (p=0.19).

**Figure 3:**
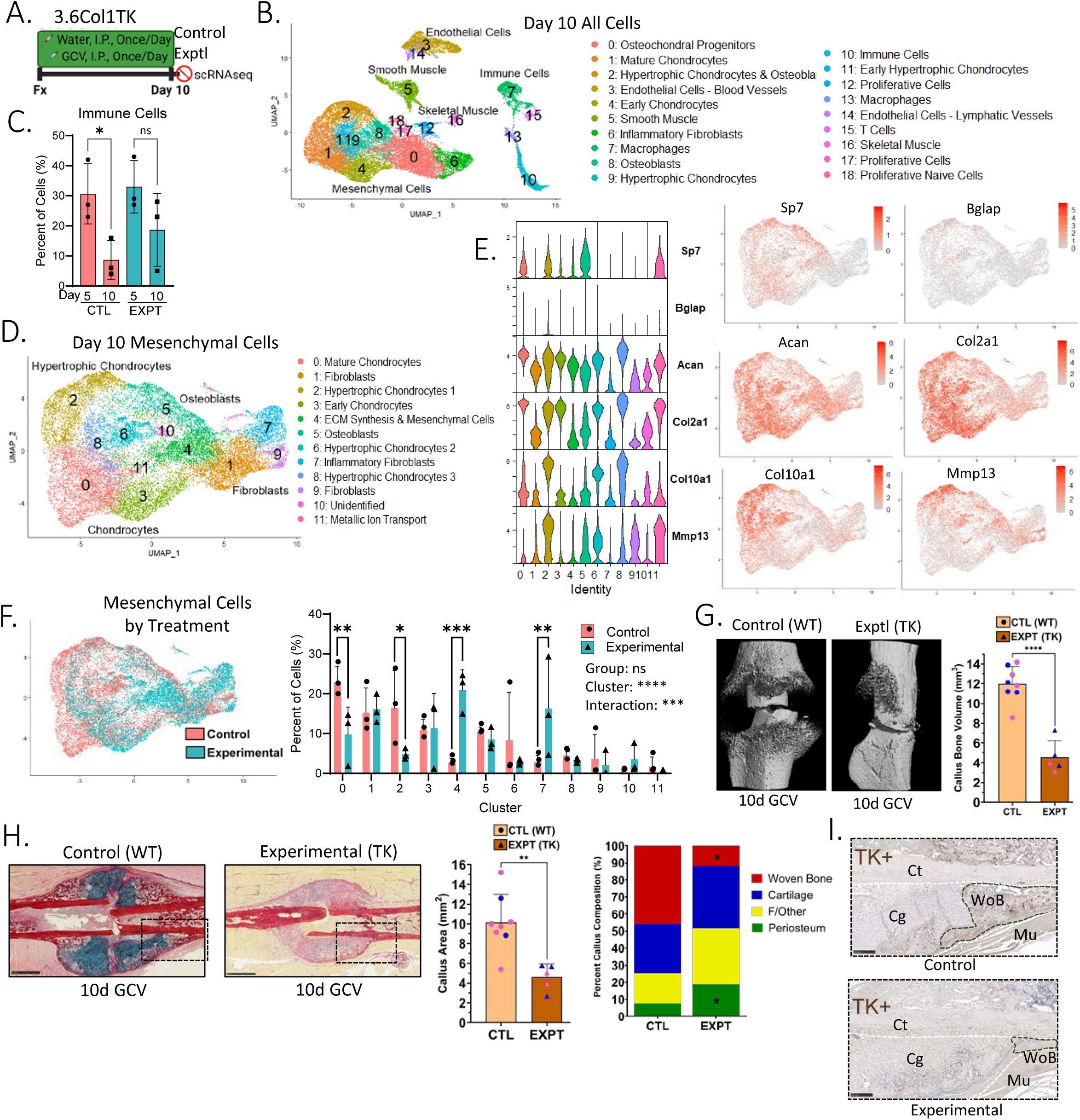
Ablation of proliferating 3.6Col1a1 lineage cells for 10 days depletes callus osteoblasts and chondrocytes and reduces callus bone volume. **(A)** After femur fracture Col1-TK mice were treated with water (Control, n=3) or GCV (Exptl, n=3) for 10 days. Cells isolated from whole-fracture callus were subjected to scRNAseq. **(B)** UMAP plot of total callus cells with annotated cluster identities. **(C)** The number of cells from immune clusters on Day 5 (Clusters 2, 6, 7, 10, 11, 12, 14, 16) and Day 10 (Clusters 7, 10, 13, 15) normalized to total cell numbers. **(D)** Mesenchymal cells were selected *in silico* and re-clustered. **(E)** Expression of canonical genes for osteoblasts, chondrocytes, and hypertrophic chondrocytes displayed as violin plots and projected on UMAPS. **(F)** Mesenchymal cells were identified by treatment group, and the percent of cells in each cluster was quantified (n=3). **(G)** Callus bone volume was quantified by microCT (n=5-8). **(H)** Histological sections of callus at 10 DPF were analyzed to determine total callus area and callus tissue composition. **(I)** Representative images of TK IHC of control (Col1-TK + water) and exptl (Col1-TK + GCV). (Statistics: two-Way ANOVA with Holm-Sidak Post Hoc test (C, F) or two-tailed t-test (G, H). Scale bars: PSRAB = 1 mm, TK=1 mm.)

To enhance resolution of the mesenchymal populations, cells in original Clusters 0, 1, 2, 4, 6, 8, 9 and 11 (Fig 3B) were re-clustered. The resulting 12 clusters were mainly comprised of different chondrocyte sub-types, with broad expression of *Acan* and *Col2a1* and more restricted expression of *Col10a1* and *Mmp13* (Fig 3D, E). Additional clusters comprised smaller populations of osteoblasts, fibroblasts, and ECM-synthesizing cells. Mature and hypertrophic chondrocytes in Clusters 0 and 2 were significantly depleted in callus from Experimental mice (15% of cells) compared to Control (39%; Fig 3F). In contrast, cells characterized by extracellular matrix (ECM) synthesis and inflammatory fibroblasts (Clusters 4 and 7; Suppl Fig S3E) were enriched in callus from Experimental mice (37% of cells) compared to Control (4%). Notably, when we selected cells based on high expression of skeletal stem and progenitor cell (SSPC) markers (*Acta2*, *Ly6a*, *Itgav*, *Thy1*, *Ctsk* (6, 21, 22)), the proportion of these cells was not different between Control and Experimental callus tissue (Suppl Fig S3D), suggesting that ablating proliferating 3.6Col1a1-lineage cells does not deplete SSPCs. Thus, scRNAseq indicates that ablation of proliferating 3.6Col1a1-lineage cells in the first 10 DPF leads to depletion of osteochondral cells, enrichment of inflammatory fibroblasts, and persistence of immune cells.

Analysis of microCT at 10 DPF revealed that callus bone volume was reduced by 60% in Experimental mice compared to Control (Fig 3G), while histological analysis revealed 50% smaller callus area (Fig 3H). Callus composition was also affected, with calluses from Experimental mice having less woven bone, but more fibrous and periosteal tissue (Suppl Fig S3E). Like 5 DPF, immunostaining for HSV-TK confirmed the loss of TK+ cells in woven bone regions of callus from Experimental (GCV-treated Col1-TK) mice compared to CTL (water-treated Col1-TK) mice (Fig 3I). Thus, scRNAseq and histological data indicate that ablation of proliferating 3.6Col1a1-lineage cells in the first 10 days after fracture depletes cartilage and bone cells in the fracture callus, with a shift toward increased fibroblasts and fibrous tissue.

### Development of a ROSA-TK mouse to drive cell-specific TK expression

Results using Col1-TK mice, in which the 3.6Col1a1 promoter drives HSV-TK expression, show an essential role for proliferation of osteogenic cells in fracture healing, which suggests that osteoblasts in this context are not post-mitotic. But a limitation of this finding is that the 3.6Col1a1 promoter is active in early osteoblasts and periosteal fibroblasts (20). To specifically target proliferating osteoblasts at different stages of differentiation, we generated a Cre-inducible mouse. A ROSA26-targeting vector was used to knock-in an expression cassette containing the HSV-TK gene downstream of a floxed stop cassette (ROSA26-LSL-HSV-TK, or ROSA-TK) (Suppl Materials; Suppl Figs S4, S5A-C). Validation of this model was done by crossing ROSA-TK mice with Osx-CreERT2 mice, which targets early and mature osteoblasts (23, 24). We confirmed tamoxifen (TMX)-inducible, Cre-mediated excision of the stop cassette and bone-specific TK expression (Suppl Fig S5D-F).

### Ablation of proliferating Osx-CreERT2-expressing cells impairs fracture callus formation, phenocopying results from Col1-TK mice

To further examine the role of proliferating early and mature osteoblasts in bone repair, we assessed fracture healing in Osx-CreERT2; ROSA-TK mice. Cre was activated 2 weeks prior to fracture by treating mice with TMX (Fig 4A). Because osterix expression promotes pre-osteoblast proliferation (25) and because Osx and 3.6Col1a1 promoters are active in similar cell types, we hypothesized that Osx-CreERT2; ROSA-TK mice would have a similar phenotype as Col1-TK mice. We first assessed HSV-TK expression by immunostaining the fracture callus of mice treated with water, rather than GCV, to allow normal healing. We confirmed that Cre-/TK+ mice do not express TK in the callus at 14 DPF, whereas Cre+/TK+ mice have strong TK expression in osteoblasts in the woven bone regions of the callus (Fig 4B) consistent findings in Osx-CreERT2 reporter mice (24). Subsequent experiments included three genotype controls (Cre-/TK-, Cre-/TK+, Cre+/TK-) in addition to experimental (Cre+/TK+) mice, all treated with TMX and GCV (except where noted). Fracture healing was evaluated 2 weeks post-fracture, when the normal callus includes both woven bone and cartilage, but little fibrous tissue.

**Figure 4:**
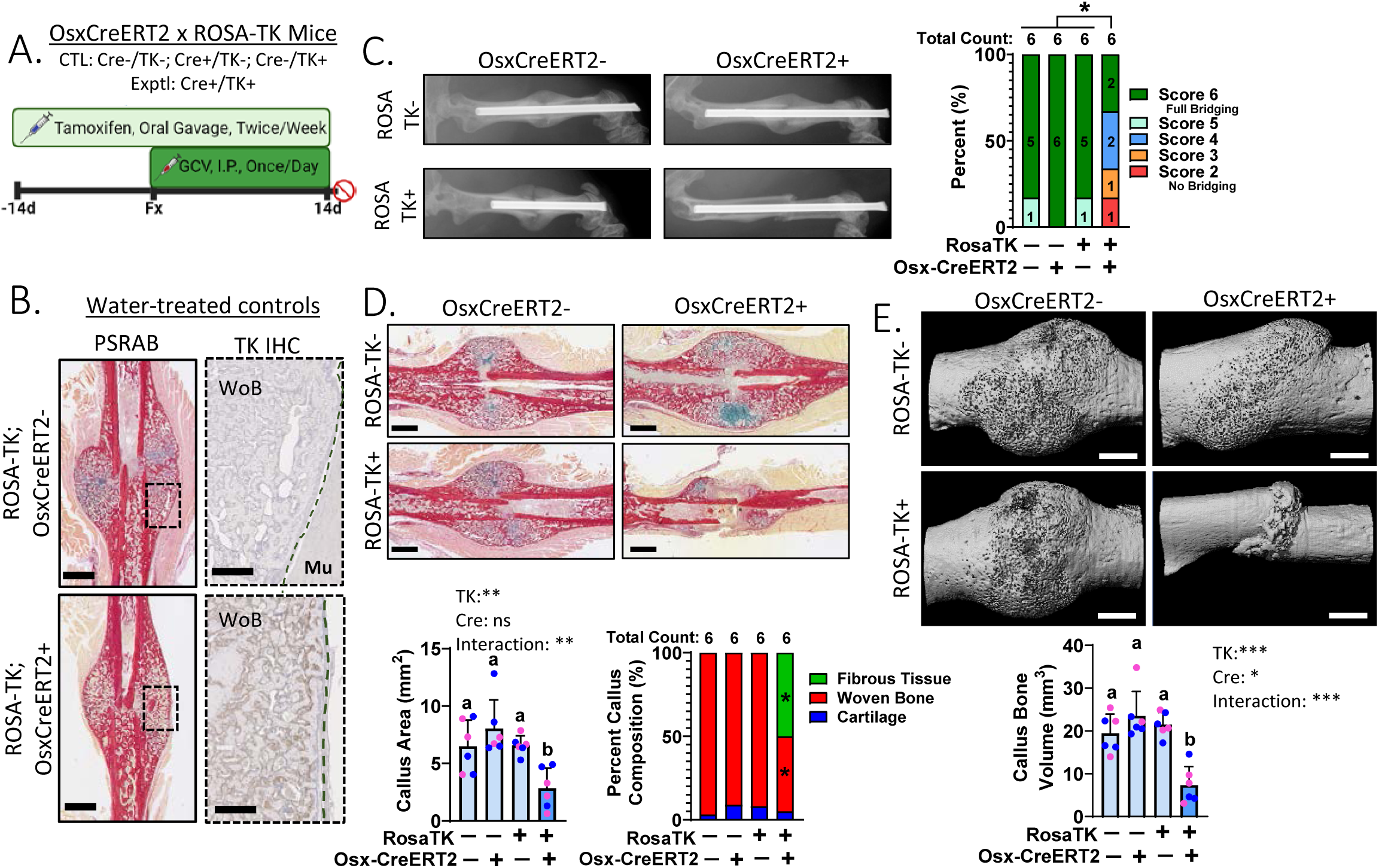
Ablation of proliferating Osterix-expressing cells impairs fracture callus bridging and mineralization. **(A)** Mice were treated with TMX starting at 10-wks age, followed by femur fracture at 12-wks, followed by 2 weeks GCV treatment. Healing was assessed 14 days post-fracture. Genotype controls were mice lacking either Osx-CreERT2 (Cre-) or ROSA-TK (TK-) alleles; experimental mice had both alleles (Cre+/TK+) (n=6). (**B)** TK+ mice were treated with water for TK IHC, which shows robust TK expression in callus woven bone only in Cre+ mice (green dashed line=callus periphery; WoB=Woven Bone; Mu=Muscle). **(C)** (Left) Representative radiographs. (Right) Quantification of radiographic healing. (**D)** Histological analysis of PSRAB stained-sections to assess callus size and composition. **(E)** Callus bone volume was quantified by microCT. (Statistical differences determined by Chi-Square test (C) or by two-way ANOVA with Holm-Sidak Post-Hoc test (D, E); *p<0.05, **p<0.01. Scale bars: TK=0.25 mm, PSRAB=1 mm, microCT=1 mm.)

Radiographic analysis indicated that 89% of Cre-control mice had a fully bridged fracture callus, whereas only 33% of Osx-CreERT2; ROSA-TK were fully bridged (Fig 4C). Histologically, fracture calluses from each control group were of similar size and were comprised predominantly of woven bone, with a small amount of cartilage and no fibrous tissue. In contrast, fracture calluses from experimental mice were significantly smaller than control, with less woven bone and more fibrous tissue (Fig 4D, Suppl Fig S6A). MicroCT analysis showed that the calluses from each control group were similarly sized, whereas experimental mice had smaller calluses with less callus bone volume (Fig 4E, Suppl Fig S6B). Measures of bone morphology at the mid-diaphysis of contralateral, intact femurs were not significantly different between control and experimental mice, indicating that short-term ablation of proliferating Osx-expressing cells does not alter bone structure (Suppl Fig S6C). Notably, significant changes in callus size and composition only occurred in Cre+/TK+ mice treated with both TMX and GCV (Suppl Fig S6D, E). In all, targeting early osteoblasts in Osx-CreERT2; ROSA-TK mice significantly impaired fracture callus bone formation and phenocopied results from Col1-TK mice. These results support the use of the ROSA-TK mouse to target proliferating cells in a Cre-dependent manner.

### Sp7/Osterix-expressing cells proliferate throughout early fracture healing

The impaired callus formation in OsxCreERT2; ROSA-TK mice indicates that proliferation of Osterix-expressing cells is essential for fracture healing. To better characterize cell proliferation during healing, cell cycle stages were assigned to the scRNAseq data using described gene markers (26). Approximately 20% of all mesenchymal cells were in the cell cycle (S or G2M phase) at both 5 and 10 DPF (Fig 5A), indicating that even 10 days after fracture, when the callus is well patterned, there is abundant cell proliferation. Of cells with high *Sp7* gene expression (≥ 90% of *Sp7* mean value), about 30% are in the cell cycle at 5 DPF, and 20% at 10 DPF (Fig 5B). Thus, based on their transcriptome, many *Sp7*-expressing cells proliferate during fracture healing.

**Figure 5:**
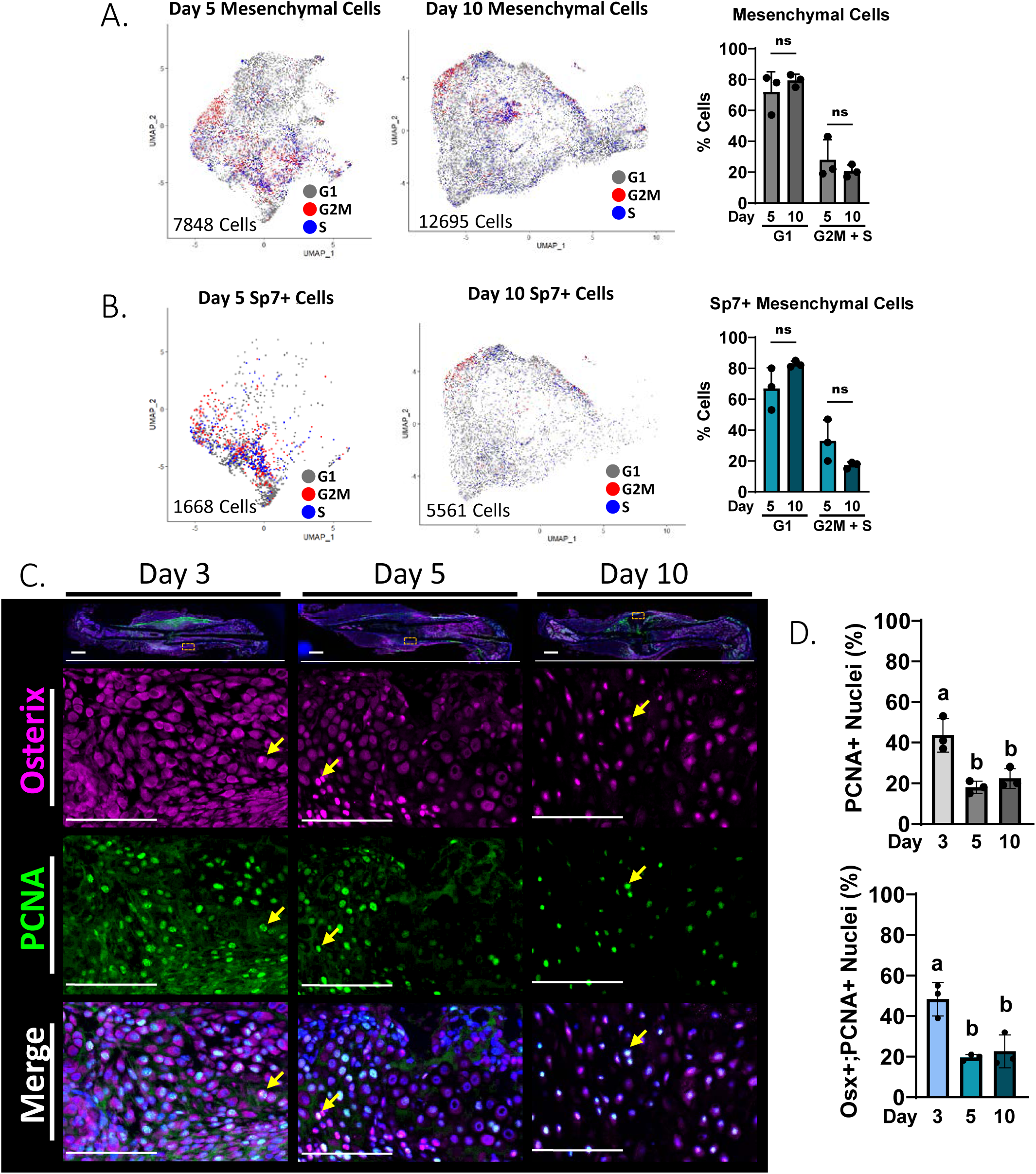
Sp7/Osterix-expressing osteoblasts proliferate during fracture healing. **(A)** Mesenchymal cells from scRNAseq analysis at days 5 and 10 post-fracture annotated for cell cycle stage. (**B)** Mesenchymal cells with high *Sp7* gene expression annotated for cell cycle stage. **(C)** Representative fracture callus sections from Control mice at 3, 5 and 10 days post-fracture stained by immunofluorescence for Osx and proliferation marker PCNA. (Scale bars: Low-Power=1 mm, Zoom=0.1 mm; Dashed yellow line=cortical bone; Dashed white line= callus periphery; Yellow Arrows=Osx+;PCNA+ cells.) **(D)** Quantification of percent PCNA+ nuclei and Osx+;PCNA+ nuclei. (n=3; graphs depict mean±SD; statistical differences determined by two-way ANOVA (A, B) or one-way ANOVA (D) with Holm-Sidak Post-Hoc.)

We next performed immunofluorescence (IF) staining of callus sections from control mice to visualize expression of PCNA (proliferating cell nuclear antigen, which is expressed during S-phase (27, 28)) and Osterix (Fig 5C). PCNA was broadly expressed in cells of the early fracture callus; approximately 40% of cells at 3 DPF had PCNA+ nuclei, while ∼20% of cells were PCNA+ at 5 and 10 DPF (Fig 5D). Notably, approximately 50% of Osterix+ cells in fracture callus were also PCNA+ at 3 DPF, and the proportion of Osterix+/PCNA+ cells remained relatively high (∼20%) at 5 and 10 DPF. Osterix+/PCNA+ cells were observed throughout the callus. Thus, by scRNAseq and IF we find evidence that a substantial proportion of proliferating cells express Osterix, consistent with the functional impairment in callus formation when these cells are ablated in Osx-CreERT2; ROSA-TK mice.

### Proliferation of Osteocalcin-lineage cells is necessary for fracture healing

As osteoblasts mature and initiate matrix mineralization, they express osteocalcin (Ocn) (29) and are said to exit the cell cycle (13). Thus, we hypothesized that there is negligible proliferation of Osteocalcin+ cells after fracture and little contribution of these cells to callus formation. To test this hypothesis, we crossed ROSA-TK mice with Ocn-Cre mice to drive TK expression in mature osteoblasts, and subjected Ocn-Cre; ROSA-TK mice and their genotype controls to femur fracture (Fig 6A). In mice treated with water (not GCV) after fracture, immunostaining confirmed that TK is strongly expressed in the woven bone callus of Cre+/TK+ mice, but not Cre-/TK+ mice (Fig 6B). We also observed some TK+ cells in the cartilage of the callus. in mice treated with GCV for 14 days after fracture, radiographic analysis showed that 77% of genotype controls had full bridging of mineralized callus, compared to only 33% of experimental mice (Fig 6C). Histologically, callus area was significantly reduced in experimental mice and callus composition was altered, with less woven bone and more fibrous tissue in Cre+/TK+ mice compared to controls (Fig 6D, Suppl Fig S7A). By microCT analysis, callus bone volume and other measures of callus morphology at 14 DPF were significantly diminished in Cre+/TK+ mice compared to controls, whereas bone morphology in non-fractured contralateral femurs was normal (Fig 6E, Suppl Fig S7B-D).

**Figure 6.**
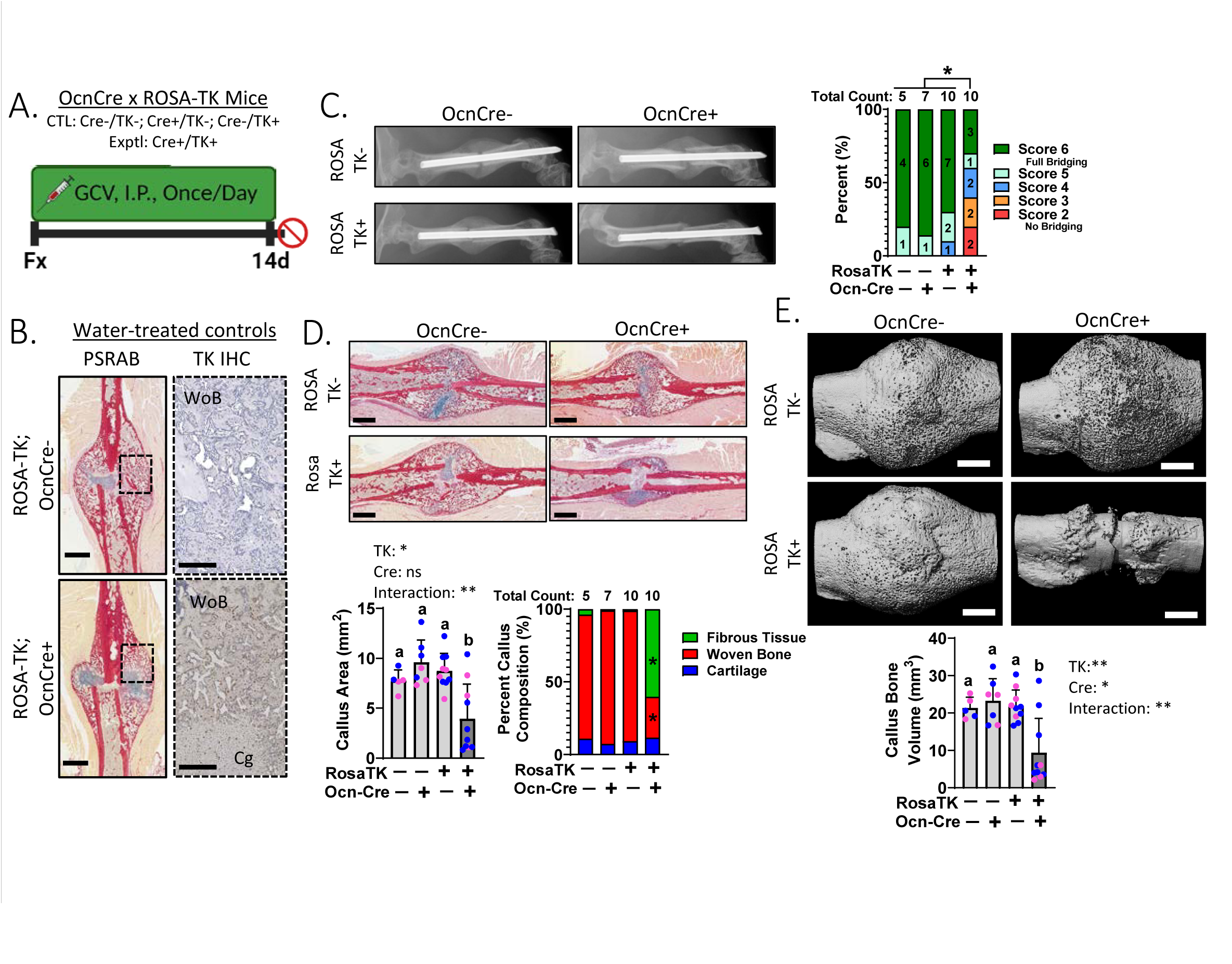
Ablation of proliferating Osteocalcin-lineage cells impairs fracture healing. **(A)** Femur fracture was performed at 12-wks age, followed by 2 weeks GCV treatment. Healing was assessed 14 days after fracture. Genotype controls included mice lacking either Ocn-Cre or RosaTK alleles; experimental mice were Cre+/TK+ (n=5-10). **(B)** TK+ mice were treated with water for HSV-TK IHC, which shows TK expression in callus woven bone only in Cre+ mice. **(C)** (Left) Representative radiographs. (Right) Quantification of radiographic healing. **(D)** Histological analysis of fracture callus composition on PSRAB-stained slides showed that experimental mice have altered composition, with less woven bone and more fibrous tissue compared to control. **(E)** Callus bone volume quantified by microCT. (Statistical differences determined by Chi-Square test (C) or two-way ANOVA with Holm-Sidak Post-Hoc test (D, E). Scale bars: TK=0.25 mm, PSRAB=1 mm, microCT=1 mm.)

To assess whether impaired healing of Ocn-Cre; ROSA-TK mice at 14 DPF could recover with time, we evaluated healing 12 weeks following fracture, i.e., 10 weeks after stopping GCV treatment. We observed radiographically and histologically that callus formation in Cre+/TK+ mice remained impaired, with incomplete bridging and persistence of fibrous tissue (Suppl Fig S8A-B). Because the impaired fracture healing of Ocn-Cre; ROSA-TK mice was unexpected, we examined healing at earlier time points. In Cre-/TK+ control mice, callus area increased progressively from 3 to 5 to 10 days, with increasing woven bone and cartilage and decreasing fibrous tissue, consistent with normal healing (Suppl Fig S8C-D). In contrast, in Cre+/TK+ experimental mice callus size and composition did not change over time. Thus, ablation of proliferating Ocn-lineage cells significantly impairs early callus formation, resulting in a fibrotic callus phenotype that persists even after withdrawal of GCV, indicating that proliferation of mature osteoblasts is required for normal fracture healing.

### Bglap/Osteocalcin expressing cells proliferate during fracture healing

To characterize the proliferative state of mature osteoblasts in fracture callus, we queried the scRNAseq data for cell cycle status of cells expressing *Bglap*, the gene that encodes osteocalcin. At 5 and 10 DPF, 15-20% of *Bglap*+ cells are in the cell cycle (Fig 7A). While the proportion of *Bglap*+ cells in the cell cycle does not change from 5 to 10 days, the total number of *Bglap*+ cells increases over 25-fold from 5 DPF (72 cells) to 10 DPF (1933 cells), indicating a large increase in the number of *Bglap*-expressing cells that are proliferating. We next visualized Osteocalcin and PCNA expression in sections of fracture callus from control mice (Fig 7B). Similar to results from OsxCre-ERT2; ROSA-TK mice (Fig 5D), at 3 DPF approximately 40% of all callus cells have PCNA+ nuclei, a rate that decreases to approximately 20% at 5 and 10 DPF (Fig 7C). Of cells that are Osteocalcin+ at 3 DPF, 40% also express PCNA; at 5 and 10 DPF 20% are OCN+/PCNA+. Thus, a substantial proportion of OCN+ cells proliferate early in fracture healing. Taken together, the functional and expression data support that Osteocalcin-expressing cells have the ability to proliferate after fracture and that this function is vital for callus formation.

**Figure 7.**
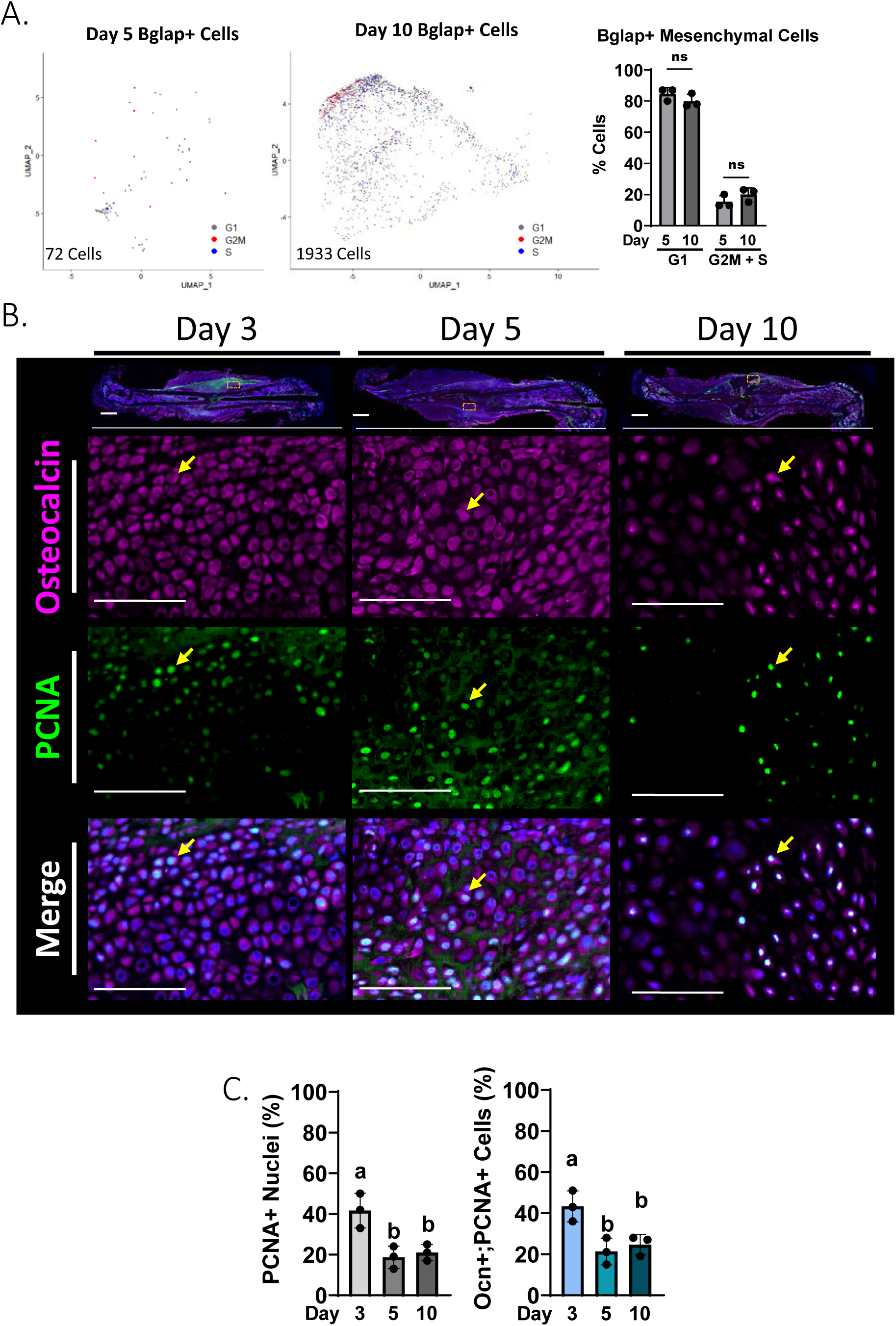
Bglap/Osteocalcin-expressing osteoblasts proliferate during fracture healing. **(A)** Mesenchymal cells from scRNAseq analysis at 5 and 10 DPF with high *Bglap* expression were annotated for cell cycle stage (n=3). **(B)** Representative callus sections from Control mice at days 3, 5 and 10 post-fracture stained by immunofluorescence for Ocn and PCNA. (Scale bars: Low-Power=1 mm, Zoom=0.1 mm; Ct.Bone=cortical bone; Callus=callus periphery; Yellow Arrows=Ocn+;PCNA+ cells.) **(C)** Quantification of percent PCNA+ nuclei and Ocn+;PCNA+ nuclei. (Graphs depict mean±SD; statistical differences determined by two-way ANOVA (A) or one-way ANOVA (C) with Holm-Sidak Post-Hoc.)

### Proliferation of Dmp1-expressing cells is necessary for fracture callus bridging and mineralization

While the Ocn-Cre mouse is widely used to target mature osteoblasts (30), a limitation is that Cre is constitutively active and thus TK will be expressed in all Ocn-lineage cells, not only cells expressing Ocn during fracture repair. In addition, Ocn-Cre may target chondrocytes during fracture healing (Fig 6B). Therefore, we crossed ROSA-TK mice with Dmp1-CreERT2 mice to induce TK expression in cells expressing the late-stage osteoblast and osteocyte marker Dmp1, at the time of fracture (Fig 8A). We confirmed strong TK expression in osteoblasts and osteocytes but not chondrocytes during fracture healing (Fig 8B), consistent with a prior report (24). Dmp1-CreERT2; ROSA-TK mice treated with GCV had significantly impaired healing compared to Cre-controls, based on radiographic, histological and microCT analyses (Fig 8C-E). Thus, ablation of proliferating Dmp1-expressing mature osteoblasts during fracture healing leads to a failure of mineralized callus formation with an increase in fibrous tissue, comparable to results seen when targeting cells using 3.6Col1a1, Osx and Ocn promoters.

**Figure 8:**
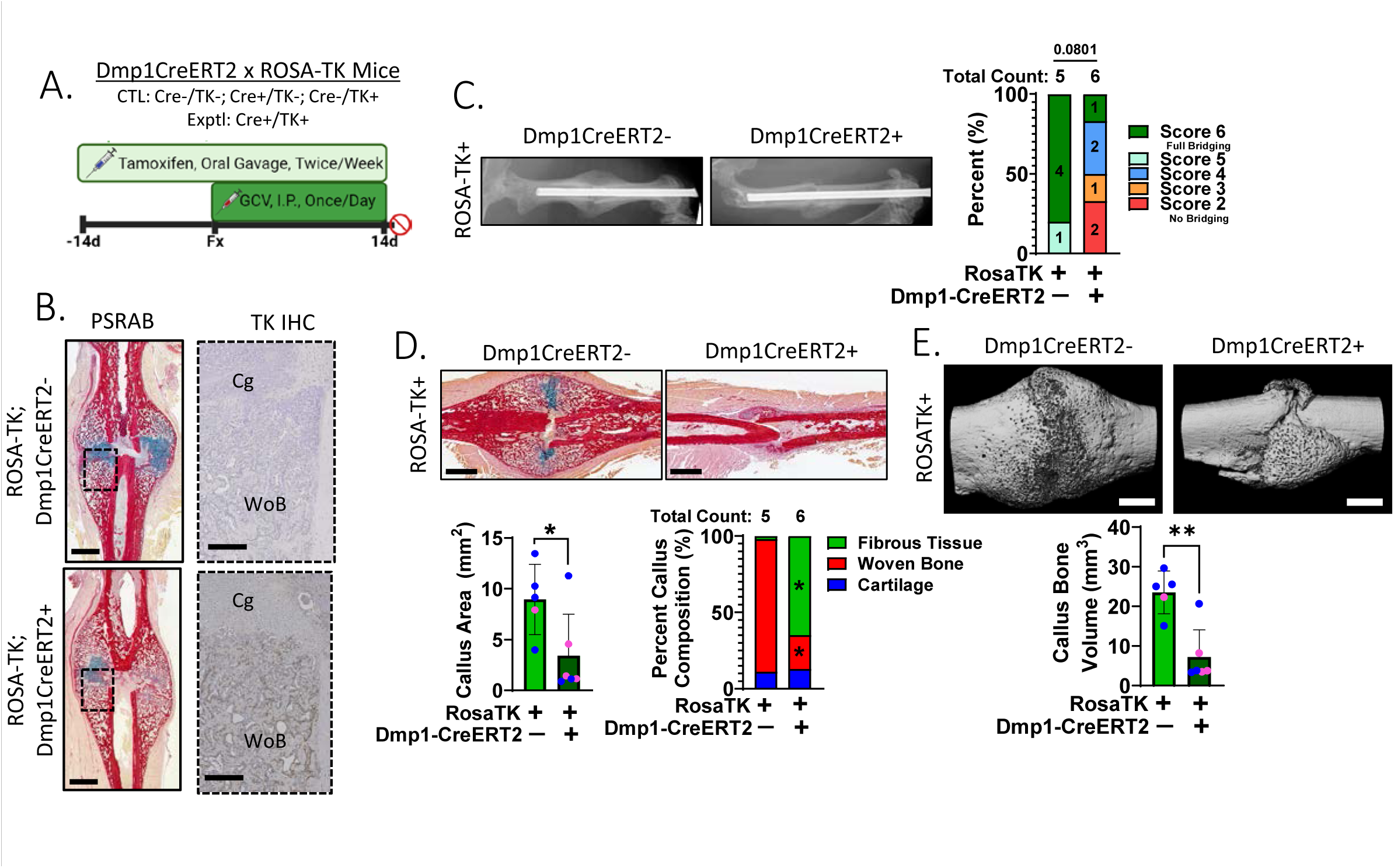
Ablation of proliferating Dmp1-expressing cells impairs fracture callus bridging and mineralization. **(A)** Mice were treated with TMX starting at 10-wks age, followed by femur fracture at 12-wks, followed by 2 weeks GCV treatment. Healing was assessed 14 days after fracture. Control mice were Cre-;TK+ while experimental mice were Cre+;TK+. **(B)** TK+ mice were treated with water for HSV-TK IHC, which shows robust TK expression in callus woven bone only in Cre+ mice. **(C)** Fracture callus bridging was quantified based on radiographic images. **(D)** Histological analysis of PSRAB stained-sections was used to assess callus size and composition. **(E)** Callus bone volume was quantified using microCT. (Statistical differences were determined by Chi-Square test (B) or by two-tailed t-test (C, D). Scale bars: PSRAB=1 mm, TK IHC=0.25 mm, microCT=1 mm.)

### Impaired fracture healing in ROSA-TK mice is not due to a gap-junction-mediated bystander effect

A feature of the TK/GCV system that has been noted in some contexts is a gap junction-mediated bystander effect, whereby TK-phosphorylated GCV can be passed from TK-expressing cells to non-TK-expressing neighbors, thus inducing apoptosis indirectly (31, 32). To test whether this effect contributes to impaired fracture healing in ROSA-TK mice, we treated Osx-CreERT2; ROSA-TK, Ocn-Cre; ROSA-TK, and Dmp1-CreERT2; ROSA-TK mice and their respective controls with the gap junction blocker Carbenoxolone (CBX) once daily (along with GCV) following femur fracture. MicroCT analysis at 14 DPF showed significantly diminished callus bone volume in Cre+/TK+ mice, regardless of CBX treatment (Suppl Fig S9 and Supplemental Results). In sum, treatment with a gap junction inhibitor did not rescue impaired fracture healing in any of the ROSA-TK models, and thus we find no evidence of a bystander effect.

### Ablation of proliferating adiponectin-expressing cells modestly impacts fracture callus formation

Because the three lines of Cre; ROSA-TK mice tested above had impaired callus formation, we asked whether there is an inherent defect in fracture healing in Cre+/ROSA-TK+ mice treated with GCV. To address this, we evaluated fracture healing in Adq-CreERT2; ROSA-TK mice (Suppl Fig S10A). Adiponectin (Adq) is expressed by marrow but not periosteal osteo-progenitors (33), and thus we hypothesized that fracture healing (predominantly a periosteal phenomenon in our model) would be minimally affected in these mice. In support of the hypothesis, we observed a modest impairment in radiographic healing and non-significant reductions in callus size and callus bone volume in Adq-Cre+/ ROSA-TK+ mice, while callus composition and BMD were unaffected (Suppl Figs S10B-D and Supplemental Results). Thus, we conclude that there is not an inherent defect in fracture healing Cre+/ROSA-TK+ mice.

## Discussion

Our results reveal a critical role for the proliferation of early and mature osteoblasts in the formation of the mineralized callus that is essential for fracture healing. Proliferation of periosteal cells is a hallmark of early callus formation (9–11), yet the identity of these proliferating cells, apart from their histological appearance, has not been well documented. Based on scRNAseq and immunofluorescence, as well as functional results whereby proliferating cells are targeted in TK-expressing mice, we find that early (3.6Col1a1+, Osx+) and mature (Ocn+, Dmp1+) osteoblasts proliferate in the first 10 days following fracture. This result is contrary to the view that mature osteoblasts are post-mitotic (7, 13, 14), a view supported by *in vitro* (12, 15) and observational *in vivo* studies of the periosteum early in skeletal development. In classic studies that focused on rapid appositional bone growth in embryonic and 2-wk old animals, cell proliferation was commonly observed in pre-osteoblasts (adjacent to but not on the bone surface), whereas proliferation of mature (surface) osteoblasts was rare (16, 17). Thus, proliferation of pre-osteoblasts, followed by their differentiation, appears to be the major source of matrix synthesizing periosteal osteoblasts early in the developing skeleton. On the other hand, studies in older mice have found evidence of osteoblast proliferation. Corral et al. (34) showed that ablation of proliferating Ocn+ osteoblasts in 6-10 wk old mice caused profound depletion of endosteal osteoblasts and reduced bone formation. Using lineage tracing in 5-mo old mice, we previously reported that up to 20% of Osx+ and Dmp1+ periosteal cells proliferate after mechanical loading and become incorporated into newly formed lamellar and woven bone (35, 36). Taken together, these findings support an important role for osteoblast proliferation as a source of bone-forming osteoblasts in the postnatal skeleton.

The finding that proliferation of mature osteoblasts is vital to the regenerative process of fracture healing is consistent with observations in other organs, wherein replication of mature tissue-resident cells contributes to regeneration after injury. Following partial liver resection, “virtually all” hepatocytes in the remaining lobes proliferate to restore liver mass within 7 days (in rodents), a process that does not normally require stem/progenitor cells (37). And following partial pancreatectomy, proliferation of mature pancreatic β-cells is the major source for new β-cells during regeneration (38). Thus, the proliferative potential of mature cells can be activated following injury, indicating that tissue regeneration does not necessarily depend solely on expansion of progenitor cells. We note that our results do not rule out the possibility that mature osteoblasts first “de-differentiate” before re-entering the cell cycle (39); additional studies of fracture healing are needed to test this idea.

There are clinical implications to our findings. Chemotherapy agents act by inhibiting cell proliferation and have been shown to impact bone healing, consistent with our findings when osteoblast proliferation is blocked (40–42). Blocking self-replication as a source of new osteoblasts resulted in a callus comprised primarily of fibrous tissue, with an appearance and function consistent with clinical nonunion (19). In patients, the causes and radiographic features of nonunion are diverse (43, 44). Nonetheless, a common histological feature in human nonunion tissue is vascularized fibrous tissue wherein fibroblast-like cells are the predominant cell type, regardless of whether the nonunion is considered atrophic or hypertrophic (3, 4). Interestingly, fibrotic nonunion is also reported in mice with systemic inflammation (45) and local radiation injury (46), suggesting that fibrous tissue is a common outcome regardless of the underlying mechanism for nonunion.

One of the early stages of fracture healing has been described as the fibrovascular phase wherein the callus contains “undefined fibrous cells” (1) or “undifferentiated mesenchymal cells” (2). With recent advances in scRNAseq, the identity and fate of fibroblastic cells in fracture callus has become clearer. Yao et al. (47) identified a population of proliferative progenitor cells that appear to represent an intermediate differentiation stage between mesenchymal progenitors and osteoblasts/chondrocytes in the early (day 5) fracture callus of normal healing mice. These cells resemble myofibroblasts, with high expression of myofibroblast marker genes including *Acta2*, *Tagln2*, *Actg1*, and *Tpm2* (47). We observed similar cells in Cluster 0 at day 5 with high expression of these same genes (Fig 2C, Suppl Fig S2D), as well as other genes associated with cell motility, ECM and wound healing (*Tnc*, *Col5a3*, *Timp1*) (48–50). Notably, the proportion of these fibroblastic cells was greater in experimental mice with impaired callus formation than in control mice (Fig 2E). Perrin et al. (51) described “injury induced fibrogenic cells” in the callus of normal healers (days 3-7), which were an intermediate stage between skeletal progenitors and osteochondral cells. Finally, Xiao et al. (52) also identified populations of fibroblastic cells that comprised a large proportion of mesenchymal cells in the early (days 4, 7) fracture callus, and reported that these cells were more abundant at day 7 in mice with systemic inflammation than in control mice. These authors suggested that starting from a common progenitor, fibroblastic cells (enriched in poor healers) and osteochondral cells (enriched in normal healers) represent alternative fates. Collectively, these studies highlight an important role for fibroblastic cells in the early fracture callus and indicate that failure to progress beyond a fibroblast-rich callus is a signature of poor healing.

A novel feature of our study is that cells were targeted based not only on promoter activity of osteoblast genes but also based on physiologic status (i.e., proliferating). This differs from other studies of fracture healing that used diphtheria toxin-based approaches to ablate cells based on promoter activity, thus targeting all Cre+ cells (53, 54). We acknowledge that ablating proliferating cells using the HSV-TK/GCV system may have limitations. In tumors, this system can target TK^−^ cells via gap junction-mediated transfer of monophosphorylated GCV from neighboring TK^+^ cells (31). However, we found no evidence that this “bystander effect” played a role in our results, as TK^+^ mice dosed with the gap junction inhibitor carbenoxolone had no rescue of their diminished callus formation (Suppl Fig S9). We also showed that Cre+/ROSA-TK+ mice do not have an inherent defect in fracture healing, as Adq-CreERT2; ROSA-TK mice had normal callus composition and only a modest reduction in callus bone volume (Suppl Fig S10).

Ono et al. (7) have highlighted the diverse origins of osteoblasts in development and regeneration, noting that osteoblasts can arise from multiple skeletal stem and progenitor cell populations (e.g., Cxcl12+, Gli1+, Acta2+, Ctsk+) located in different tissues (e.g., marrow, growth plate, periosteum) (6, 22, 55, 56). This view has been informed by a rich literature of lineage-tracing studies, and supports a model whereby expansion of local SSPCs, followed by their differentiation is the predominant pipeline of bone-forming osteoblasts. However, the possibility that mature skeletal cells can be activated to contribute to bone regeneration has also been postulated (39). Our findings support the latter, with evidence that, in the context of fracture healing, the proliferation of early and mature osteoblasts is an essential, local source for bone-forming osteoblasts.

In conclusion, our findings indicate that proliferation of early and mature osteoblasts is essential for the formation of mineralized callus necessary for fracture healing. Blockade of osteoblast proliferation leads to depletion of callus osteoblasts and chondrocytes, and persistence of immune cells and fibroblasts. These results indicate that proliferation of mature osteoblasts can be an important source of bone-forming osteoblasts in the adult skeleton, which can be considered in new approaches to rescue fracture nonunion.

## Materials and Methods

### Chemicals and Reagents

See Supplemental Methods.

### Mouse Care

This study was approved by the Washington University IACUC. Mice were housed in standard conditions under 12 hr light/dark cycles, up to 5 per cage, with *ad libitum* food and water. Male and female mice were used in approximate equal number. Mice were euthanized via CO_2_ asphyxiation and secondary cervical dislocation.

### Col1-TK Mice

Transgenic 3.6Col1a1-TK (Col1-TK) mice were used to target proliferating (pre)osteoblasts, as described(18, 19). Mice were administered ganciclovir (GCV) starting on the day of fracture (at 12 wks age) and were euthanized at 5-, 10-, or 21-days post-fracture.

### Fracture Model

A model of mid-diaphyseal femur fracture with intramedullary pin stabilization was used (19, 57). This model heals via secondary fracture healing, with intramembranous bone formation at the periphery and endochondral bone formation within the fracture gap.

### Drug Treatments

Osx-CreERT2, Dmp1-CreERT2, and Adq-CreERT2 mice were dosed with tamoxifen (TMX, 100 mg/kg in corn oil, oral gavage, 2/wk) starting 2 wks pre-through 2 wks post-fracture. Except where noted, mice were injected with GCV (8 mg/kg in water, 1/day, IP) from the day of fracture to the indicated time (3, 5, 7, 10 or 14 days post-fracture). Some control mice were injected with either corn oil (control for TMX), or water (control for GCV). CBX (50 mg/kg, in saline, 1/day, IP) was administered from the day of fracture until euthanasia 2 wks post-fracture.

### Single Cell RNA Sequencing

For scRNAseq, 12 Col1-TK mice were fractured and treated with water (Control) or GCV (Exptl) for 5 or 10 days (n=3/time/treatment). Callus tissue was dissected, enzymatically digested, and single-cell suspensions (1/mouse) were processed for library preparation (10X Genomics). An average of 8700 cells/sample were sequenced (Illumina) resulting in detection of 23,140 genes/sample (avg.). Using Seurat (58), all cells from each time point were subjected to unbiased clustering. Based on bioinformatic analysis of top 50 DEGs per cluster as well as gene expression plots, each cluster was annotated. Clusters primarily comprised of mesenchymal cells were re-clustered for higher resolution. Cell cycle stage was determined based on canonical gene expression profiles as described (26); cells assigned to either the S or G2M phase were judged to be actively proliferating.

### Generation, Breeding and Validation of ROSA-TK Mouse Line

We generated a novel Cre-inducible mouse (ROSA26-LSL-HSV-TK, or ‘ROSA-TK’) for targeted ablation of proliferating cells. A plasmid comprising the truncated herpes simplex virus thymidine kinase allele (HSV-deltaTK)(59) downstream of a CAG promoter and floxed stop cassette (Suppl Fig S4) was knocked into the ROSA26 locus. Genomic PCR confirmed knock-in in six founder mice and F1 transmission (Suppl Fig S5A-C). Two heterozygous founders were used to establish the ROSA-TK lines backcrossed on a C57Bl/6 background. ROSA-TK mice were crossed with Osx-CreERT2 (23), Dmp1-CreERT2 (60), Ocn-Cre (Jax), and Adq-CreERT2 (Jax) mice to generate control (Cre-;TK-, Cre-;TK+, Cre+;TK-) and experimental (Cre+;TK+) mice.

### Radiographic Evaluation

Fracture healing was assessed from a lateral radiograph (Faxitron) using the modified RUST method adapted for mouse femurs (61). Scores ranged from 2 (no callus on either cortex) to 6 (fully bridged callus both cortices, but fracture line still visible).

### Micro-Computed Tomography

Fractured and intact contralateral femurs were microCT scanned (Scanco, 10.5 μm voxel) for analysis of a 600-slice ROI centered at the fracture site. A single threshold (200 per mille) was used to determine standard bone measures (BV, TV, BV/TV, vBMD, TMD), inclusive of callus and cortical bone. To exclude the cortical bone within the ROI, a two-threshold method was used (lower=150, upper= 460 per mille), from which callus bone volume was determined.

### Histological Analysis of Callus

Femurs were decalcified and embedded in paraffin. Sagittal sections, 5 μm thick, were cut and stained by H&E and picrosirius red/alcian blue (PSRAB). Slides were scanned and the periosteal callus was analyzed to determine % area of cartilage, woven bone, fibrous tissue and periosteum (Suppl Fig S1C).

### Thymidine Kinase (TK) Immunohistochemistry

Immunohistochemistry was used to visualize HSV-TK expression in the callus region of histological sections, as described (62).

### Co-Immunofluorescence

To visualize proliferating osteoblasts, histological sections were co-stained with either anti-PCNA (1:500) and anti-Osterix (1:150), or anti-PCNA and anti-Osteocalcin (1:500) and visualized by fluorescence microscopy. Proliferating cell nuclear antigen (PCNA) is expressed during the S phase of the cell cycle and its staining in fracture callus closely mimics BrdU incorporation (27, 28). QuPATH (63) was used to identify cells that were PCNA+ and either Osx+ or Ocn+. Percent of proliferating early-osteoblasts were determined as the number of PCNA+;Osx+ cells divided by the total number of Osx+ nuclei, while % of proliferating mature-osteoblasts was the number of PCNA+;Ocn+ cells divided by number of Ocn+ cells.

### Statistics

Group sample sizes were determined *a priori* based on prior results and expected effect sizes. Personnel were blinded to animal genotype and treatment during data collection and analysis. Analyses were carried out in GraphPad Prism 9.0. Statistical tests used are stated in figure captions. p < 0.05 was defined as significance.

## Acknowledgements

Supported by DOD PW81XWH-21-1-0386; NIH R21AR076636, R01 AR079246, T32 AR060719, P30 AR074992, S10 RR023660. We acknowledge the Washington University Musculoskeletal Research Center (P30 AR074992) for imaging and histology support; the Genome Technology Access Center (GTAC@MGI) for scRNAseq and COMPBIO support; the Center for Regenerative Medicine for Bioinformatic support; and the Alafi Neuroimaging Core for slide scanning (S10 RR027552). Thanks to Crystal Idleburg and Samantha Coleman for expert histology. Osx-CreERT2 mice were derived from breeders gifted from Dr. Henry Kronenberg (MGH). Dmp1-CreERT2 were derived from breeders gifted by Dr. Paola Divieti Pajevic (BU Dental School). The TALEN kit used for TALE assembly was a gift from Keith Joung. Anti-HSV-TK antibody was a gift from Dr. William Summers (Yale). The views expressed are those of the authors and do not necessarily represent the official views of the National Institutes of Health.

## Funding Acknowledgement

DOD PW81XWH-21-1-0386 (to MJS); NIH R21 AR076636 (MJS), R01 AR079246 (DMO), T32 AR060719 (AFC, KRH), P30 AR074992 (MJS), S10 RR023660 (MJS). The views expressed are those of the authors and do not necessarily represent the official views of the National Institutes of Health.

## Gould et al, Supplemental Material

### Supplemental Results

#### Blocking gap junction-mediated intercellular communication (GJIC) does not rescue impaired fracture healing in Osx-CreERT2, Ocn-Cre, or Dmp1-CreERT2; ROSA-TK mice

A feature of the TK/GCV model that has been noted is a bystander effect (1, 2). This occurs when a TK-expressing cell is linked to other cells via gap junctions, which facilitates the passage of the toxic mono-phosphorylated GCV (mediated only by TK enzymatic activity) to neighboring cells that may not express TK. This modified GCV can then be further phosphorylated and lead to apoptosis in a proliferating cell that never expressed TK. To test whether this by-stander effect contributes to the impaired fracture healing seen in ROSA-TK mice, we treated Osx-CreERT2; ROSA-TK, Ocn-Cre; ROSA-TK, and Dmp1-CreERT2; ROSA-TK mice and their respective controls with the gap junction blocker Carbenoxolone (CBX) once daily (along with GCV) following full femur fracture. Based on microCT analysis at 14 DPF, Cre+/TK+ mice had significantly diminished callus bone volume compared to Cre-/TK+ control mice, and this was not different in CBX-treated mice than in mice that were not treated with CBX (Suppl Fig S9). Thus, inhibiting gap junction communication did not rescue impaired fracture healing in any of the ROSA-TK models. This result indicates that the impaired healing observed when early and mature proliferating osteoblasts are ablated is not due to a gap junction-mediated intercellular bystander effect but is due to the ablation of actively proliferating Osterix-, Osteocalcin-, or Dmp1-expressing cells during fracture healing.

#### Ablation of proliferating Adq-expressing cells modestly affects fracture healing

Fracture callus is predominantly derived from periosteal skeletal progenitors, albeit with some contribution from bone marrow cells (3). Adiponectin (Adq) is expressed in osteoprogenitor cells in bone marrow, but is not expressed by periosteal cells (4). Therefore, we evaluated fracture healing in Adq-CreERT2; ROSA-TK mice as a negative control, i.e., with the hypothesis that fracture healing would be minimally affected. To activate Cre, mice were dosed with TMX 2 weeks prior to fracture. Cre-/TK+ (control) and Cre+/TK+ (exptl) mice were treated with TMX and GCV and fracture healing was evaluated at 14 days post-fracture (Suppl Fig 10A).

Radiographic analysis indicated that 80% of control mice had callus bridging of both cortices and 90% had bridged at least one cortex, compared to 38% and 75%, respectively, in Cre+/TK+ mice, suggesting delayed bridging in the latter (Suppl Fig 10B). Histologically, fracture calluses from Cre+/TK+ mice were marginally smaller than Cre-/TK+ controls (-25%, p=0.11) although callus composition was proportionately normal in Cre+/TK+ mice (Suppl Fig 10C). MicroCT analysis showed that callus bone volume was marginally less in Cre+/TK+ mice than in Cre-controls (-20%, p=0.14), while callus BMD was not affected (Suppl Fig 10D). In comparison, Osx-CreERT2;ROSA-TK mice have 65% lower callus bone volume compared to their controls (p<0.0001; Fig 4E). These results indicate a modest effect on fracture callus formation in mice wherein proliferating adiponectin-expressing cells were ablated, consistent with targeting of bone marrow skeletal progenitors but not periosteal progenitors. This finding supports that Cre+/TK+ mice treated with GCV do not inherently have impaired fracture healing.

### Supplemental Methods

#### Chemicals and Reagents

Tamoxifen (TMX, T5648), corn oil (C8267), and Ganciclovir (GCV, G2536) were from Millipore Sigma. Goat Anti-Rabbit Immunoglobulins/HRP (affinity isolated, P0448) was from Agilent Dako. Diaminobenzidine (DAB) chromogen (SK-4105) and Hematoxylin QS Counterstain (H-3404) were from Vector Laboratories. ReadyProbes Mouse-on-Mouse IgG Blocking Solution (R37621), Goat anti-Mouse IgG (H+L), Superclonal™ Recombinant Secondary Antibody, Alexa Fluor™ 488 (A28175), Goat anti-Rabbit IgG (H+L) Highly Cross-Adsorbed Secondary Antibody, Alexa Fluor™ 546 (A11035), and ProLong™ Glass Antifade Mountant with NucBlue™ Stain (P36981) were from ThermoFisher. Anti-Osteocalcin (A93876) and Anti-Osterix (22552) were from Abcam. Anti-PCNA (2586) was from Cell Signaling Technology. 22μm sterile filters (09-720-004) were from Fisher Scientific. Collagenase Type II (L5004176) was from Worthington. Pronase (53702), Trypan blue (T8154), and High Glucose DMEM (D6429) were from Sigma. LIVE/DEAD™ Viability/Cytotoxicity Kit (L3224) was from Invitrogen. 40μm SureStrain™ Premium Cell Strainers (C4040) were from MTCBio. MS Columns (130-042-201) were from Miltenyi Biotec. The Corded Dremel (5EEU5) was from Grainger. The Dremel bit (2901A247) and the tungsten guide wire (3775K37) were from McMaster Carr. Hypodermic tubing (304H24TW) was from Microgroup. 4-0 nylon sutures (1854G) were from Ethicon. Anti-Thymidine Kinase Antibody was a gift from William Summers (Yale).

#### Col1-TK Mice

Breeders were rederived from cryopreserved embryos gifted by Drs. O’Brien and Jilka (5) (UAMS). A total of 90 mice, age 12-wks were used. Transgenic 3.6Col1a1-TK (Col1-TK) mice were used to target proliferating (pre)osteoblasts, wherein expression of herpes simplex virus thymidine kinase (TK) in early osteoblasts is driven by the 3.6 kilobase rat *Col1a1* promoter (5, 6). Heterozygous female Col1-TK mice were bred to wild-type males to generate wild-type (WT) controls or Col1-TK (TK) experimental mice of both sexes. Mice were administered ganciclovir (GCV, 8 mg/kg i.p., 1/d) starting on the day of fracture and were euthanized at 5-, 10-, or 21-days post-fracture.

#### Fracture Model

An established model of mid-diaphyseal femur fracture was used (6, 7). Briefly, following administration of an analgesic (Buprenorphine SR, 1 mg/kg, s.c), mice were anesthetized (1-3% isofluorane) and a 5 mm incision was made above the knee, exposing the distal femoral condyles. The medullary canal was reamed out, and a 1.5-inch-long tungsten guide wire was inserted into the marrow space. To create a mid-point femur fracture, a transverse force was applied using a three-point bending set-up (DynaMight 8841; Instron). A 24-gauge, 1-inch hypodermic stainless-steel tube was inserted over the guidewire to stabilize the fracture. The guidewire was removed, the tube clipped to length, and the incision closed using 4-0 nylon suture. Pin placement was confirmed with radiography immediately after surgery. This model heals via secondary fracture healing, with intramembranous bone formation at the periphery and endochondral bone formation within the fracture gap.

#### Cell Isolation for Single Cell RNA Sequencing

For scRNAseq, Col1-TK mice (n=3/time/treatment) were fractured and treated with either water (Control) or GCV (Exptl) for 5- or 10-days post-fracture. Following euthanasia, the fracture site was exposed, and total callus tissue scraped away from cortical bone using a fresh scalpel. Tissue was minced in 1 mL of 22 mm sterile-filtered digestion buffer comprised of 1663 U/mL Collagenase Type II and 1814 PUK/mL Pronase in High Glucose DMEM with 8% FBS. After digestion for 1 hr at 37°C, cells were pelleted and re-suspended in 0.01% BSA in sterile PBS. Cells were filtered through a 40 mm filter, and dead cells were removed using an MS Column. Trypan Blue-stained cells were counted and viability estimated using a Live/Dead kit. Twelve samples (one per mouse, no pooling) comprising 40,000 cells/sample with 70% viability were sent for sequencing.

#### Single Cell RNA Sequencing and Analysis

Twelve single cell suspensions were transferred to the Genome Technology Access Center at Washington University for library preparation using the 10X Genomics Chromium Single Cell 3’ Reagent Kit (v3.1). Sequencing was done in a single lane of Illumina Novaseq (avg. 8,700 cells per sample; range: 4170-16,323). A mean of 74,692 reads per cell were sequenced, and mapped to a median of 2,770 genes per cell, with overall coverage of 61% of the transcriptome. An average of 23,140 genes were detected per sample. Following standard normalization and filtering, data from 104,740 cells (Day 5: Control 32,322 cells; GCV 28,383; Day 10: Control 23,278; GCV: 20,307) were available for analysis.

Secondary analysis was done using Seurat Version 4.4.0 in R (8). All cells from each time point, independent of treatment, were subjected to unbiased clustering and were visualized on UMAP plots. For cluster annotation, the top 50 differentially expressed genes (DEGs) from each cluster were entered into Comprehensive Multi-Omics Platform for Biological Interpretation (COMPBIO, Washington University) to identify the top biological themes and components for each cluster. Following annotation, clusters comprised of predominantly mesenchymal lineage cells were digitally selected and re-clustered for better cell identity resolution. The top 50 DEGs from the mesenchymal-only clusters were also input into COMPBIO for annotation. Seurat in R was used for cell subsetting. Threshold cutoffs for *Sp7*/Osterix and *Bglap*/Osteocalcin used were 90% of the mean gene expression of mesenchymal cells only from control water-treated mice from each time point. Cell cycle stage was determined based on canonical gene expression profiles as described (9). Cells are assigned to either G1 phase (stagnant, functional phase), S Phase (actively replicating DNA in preparation for cell division), or G2M phase (preparing for and going through mitosis). Cells assigned to either the S or G2M phase are judged to be actively proliferating.

#### Generation and Validation of a ROSA-TK Mouse Line

To generate a Cre-inducible mouse (ROSA26-LSL-HSV-TK, or ‘ROSA-TK’) for targeted ablation of proliferating cells, we utilized a truncated herpes simplex virus thymidine kinase allele (HSV-deltaTK, Addgene 114273) to circumvent male sterility caused by the full sequence TK while retaining enzymatic function(10). The HSV-deltaTK knock-in targeting vector contained a CAG promoter, followed by floxed stop cassette (LSL), the HSV-deltaTK allele, a P2A self-cleaving site, and finally an mCherry allele (Suppl Fig S4). The plasmid was synthesized by the Genome Engineering and Stem Cell Core (Washington University). The floxed stop cassette allows for tissue-specific expression of HSV-deltaTK and mCherry upon Cre-mediated excision. The P2A site allows for bicistronic expression of the HSV-deltaTK and mCherry genes. ROSA26 specific TALENs were designed using the ZiFit targeter (http://zifit.partners.org) and ROSA26 TALENs binding sites as follows:

1) 5’ ROSA26 TALEN binding site: 5’ TCCCTCGTGATCTGCAACTCC 3’
2) 3’ ROSA26 TALEN binding site: 5’ GGGCGGGAGTCTTCTGGGCA 3’

The TALEN kit used for TALE assembly was a gift from Keith Joung (Addgene kit # 1000000017). DNA fragments encoding ROSA TALEN repeat arrays were cloned into plasmid pJDS71. ROSA TALENs plasmids were linearized for *in vitro* transcription with EcoRI and TALENs RNA was synthesized using the mMessage mMachine T7 Ultra kit (Ambion) and purified with Megaclear columns (Ambion). The knock-in cassette was introduced into the mouse genome via pronuclear injection of *in vitro* transcribed TALENs RNA and ROSA donor DNA into C57BL/6J;CBA hybrid oocytes which were then implanted into hosts dams (Mouse Genetics Core, Washington University). Six offspring with correctly targeted alleles were identified by long-range PCR using ROSA specific primers external to the homology arms (not shown). These six potential founders (F0 generation) were bred to WT C57Bl/6 mice. Presence of the mutant allele was demonstrated in the six potential founders (F0) using standard PCR (Suppl Fig S5A-B). F1 pups were screened to confirm transmission of the mutant allele from founders to offspring (Suppl Fig S5C).

Two confirmed ROSA-TK founders (354-7 and 356-4) were crossed to tamoxifen-inducible Osx-CreERT2 mice and validation was assessed in F1 mice by three methods. First, we confirmed tamoxifen-dependent excision of the stop cassette only in Cre+ mice (Suppl Fig S5D-E). Second, we used IHC to confirm Cre-dependent TK protein expression in trabecular osteoblasts only in Osx-CreERT2+; ROSA-TK+ mice, supporting that excision of the stop cassette is sufficient to drive TK expression in the targeted cells (Suppl Fig S5F). Third, we demonstrated nonunion fractures only in Cre+/TK+ mice (Fig 4, Suppl Fig 6). These two founders were used to establish the ROSA-TK lines, and all mice for subsequent studies were derived therefrom. Each founder line was kept separate to ensure reproducibility of phenotypes independent of founder. (Note that we did not detect mCherry fluorescence on frozen sections, but confirmed mCherry expression using IHC (not shown).)

#### Maintenance of Cre Lines and Generation of Cre;TK Mice

Osx-CreERT2 (from Dr. Kronenberg(11), MGI:4829803), Dmp1-CreERT2 (from Dr. Divieti Pajevic(12), MGI:5792965) and Ocn-Cre (Jackson Laboratory, #19509) mice were acquired and maintained by breeding Cre-positive males to wild type C57Bl/6 females (Jackson Laboratory, #000664). Cre-positive males were bred with ROSA-TK females to generate control (Cre-;TK-, Cre-;TK+, Cre+;TK-) and experimental (Cre+;TK+) littermates. When possible, Cre+;TK+ males were bred to Cre-;TK+ females to generate control and experimental littermates with homozygous TK+/+ genotypes.

#### Genotyping

For 3.6ColTK mice, genotyping was done by Transnetyx using toe biopsies for the absence or presence of the puromycin selection cassette:

- Fwd: 5’ –GCGGTGTTCGCCGAGAT– 3’; Rev: 5’ –GAGGCCTTCCATCTGTTGCT– 3’ For ROSA-TK mice, primers targeting the wild-type ROSA allele or the CAG promoter were used to distinguish wild-type and mutant (TK+) alleles:
- Wild-type: For: 5’ – GTTATCAGTAAGGGAGCTGCAGTGGAGTAG – 3’; Rev: 5’ – CCGAAAATCTGTGGGAAGTCTTGTCCCTCC – 3’
- TK+: For: 5’ - GTTATCAGTAAGGGAGCTGCAGTGGAGTAG - 3’; Rev: 5’ - CTCCACCCATTGACGTCAATGGAAAGTCCC - 3’

Following establishment of the ROSA-TK line, genotyping was done by Transnetyx using toe biopsies for the wild-type ROSA allele or the inclusion of the mCherry (TK+) sequence:

- Wild-type: For: 5’ – TTCCCTCGTGATCTGCAACTC – 3’; Rev: 5’ – CTTTAAGCCTGCCCAGAAGACT – 3’
- mCherry: For: 5’ – AGCGCGTGATGAACTTCGA – 3’; Rev: 5’ – GCGCAGCTTCACCTTGTAGAT – 3’

For Stop cassette genotyping in Osx-CreERT2; ROSA-TK mice, mice were given three doses of TMX across 5 days. Tail snips were taken prior to and after full TMX treatment for genotyping to visualize excision of the stop cassette:

- Fwd: 5’ – GAGGGCCTTCGTGCGTC – 3’; Rev – 5’ CATCGGCTCGGGTACGTAGA – 3’

For Cre allele genotyping, genotyping was done by Transnetyx using toe biopsies:

- Osx-CreERT2 and Dmp1-CreERT2: Fwd: 5’ - TGCGCCTGCTGGAAGAT -3’; Rev: 5’ – GGTTGGCAGCTCTCATGTCT – 3’
- Ocn-Cre: Fwd: 5’ – TTAATCCATATTGGCAGAACGAAAACG – 3’; Rev: 5’ – CAGGCTAAGTGCCTTCTCTACA – 3’

#### Radiographic Evaluation

Lateral radiographs were taken weekly to confirm proper positioning of the intramedullary pin and to monitor healing (Faxitron UltraFocus100). Radiographs from 14 or 21 weeks post-fracture were blindly scored using the modified RUST (radiographic union score for tibial fracture) method adapted for mouse femurs(13). From the lateral projection, we scored the anterior and posterior cortices on a 4-point scale (1=fracture line visible, no callus formation; 2=fracture line visible, visible callus; 3=fracture line visible, callus fully bridged; 4=fracture line not visible, callus fully bridged) and summed the two scores (possible range 2 to 8). Because the fracture lines were still visible on both cortices at these timepoints, the maximum score in our study was 6. Mice were excluded if the fixation pin displaced during healing.

#### Micro-Computed Tomography

Fractured and intact contralateral femurs were fixed in 10% non-buffered formalin for 16 hours, rinsed with PBS and stored in 70% ethanol. Femurs were scanned *ex vivo* using microCT (VivaCT 40, Scanco, 10.5μm voxel size, 55kV, 145μA, 300ms integration time). Conventional analysis of a 600-slice ROI centered on the mid-point of the fracture line was done, as described (6). A single lower threshold (200 per mille, contour peel 2) was used to determine bone volume (BV, mm^3^), tissue volume (TV, mm^3^), volumetric bone mineral density (vBMD, mg HA/cm^3^), and tissue mineral density (TMD, mg HA/cm^3^), inclusive of callus bone and cortical bone. Cortical parameters from the mid-diaphysis of intact limbs were similarly analyzed; the population mean (dotted line) and standard deviation (grey shading) are shown overlaying data from fractured limbs. To isolate callus bone in fractured femurs, a two-threshold method was used: 150 and 460 per mille (contour peel of 0) to segment total bone and cortical bone, respectively. Cortical bone volume (CBV_460_) was subtracted from total bone volume (TBV_150_) to yield callus bone volume (mm^3^). Following microCT, femurs were decalcified and processed for histology.

#### Histology and Analysis of Callus Composition

Femurs were decalcified in 14% EDTA, pH 7.0 for 2 weeks, with EDTA changed every 3-4 days. Femurs were embedded in paraffin wax and 5 μm thick, longitudinal sagittal sections through the middle of the fracture callus were mounted on glass slides. Slides were deparaffinized in xylenes, rehydrated in graded ethanols and stained by hematoxylin and eosin (H&E), and picrosirius red and alcian blue (PSRAB) to visualize collagen-rich woven bone and proteoglycan-rich cartilage, respectively. Slides were imaged at 20× on a Nanozoomer slide scanner (Hamamatsu Photonics). Callus composition was quantified in NDP View 2 software (Hamamatsu) by drawing custom contours around different tissue components (see example in Suppl Fig S1C). Total callus area was the area of both sides of the periosteal callus, excluding any intact cortical bone or bone marrow. Cartilage was identified by Alcian blue stained areas and woven bone was identified by Picrosirius red stained areas. Fibrous tissue was identified as a mix of the two stains that did not morphologically resemble either cartilage or bone.

#### Thymidine Kinase (TK) Immunohistochemistry

Immunohistochemistry was used to visualize HSV-TK expression in the callus region of histological sections. Following deparaffinization, endogenous peroxidase activity was blocked using 3% hydrogen peroxide. Sections were blocked with 10% goat sera + 1% bovine serum albumin for 1 hr at room temperature (RT). Anti-TK antibody (1:1000, gift from William Summers(14)) was diluted in 2% goat sera and incubated overnight at 4C. Anti-rabbit HRP (1:500) was diluted in 2% goat sera and incubated for 1 hr at RT. All samples were incubated with diaminobenzidine (DAB) chromogen for 30 sec and rinsed in water to stop the reaction. Sections were counterstained with hematoxylin for 30 sec and imaged at 20× as above.

#### Co-Immunofluorescence

Antigen retrieval was done with boiling 10mM citrate buffer and overnight incubation at 60C. Sections were permeabilized with 0.3% Triton-X 100, blocked with 10% goat serum for 1 hr at RT, and endogenous mouse IgG was blocked with Mouse-on-Mouse IgG Blocking Solution for 1 hr at RT. Sections were stained with either anti-PCNA (1:500) and anti-Osterix (1:150) or Anti-PCNA and anti-Osteocalcin (1:500) diluted in 1.5% goat serum overnight at 4C. 1:200 anti-mouse 488 (A28175) and 1:200 anti-rabbit 546 were diluted in 1.5% goat serum and incubated at RT for 1.5 hr. Sections were mounted and coverslipped with Prolong Glass with DAPI. PCNA is expressed during the S phase of the cell cycle where it interacts with DNA polymerase complexes to increase processivity by holding DNA polymerase onto the DNA strand (15). When compared to the incorporation of BrdU in rat fracture model as a marker for DNA synthesis and cell proliferation, PCNA staining closely mimicked BrdU incorporation, supporting its use in detecting proliferating cells (16).

#### Co-Immunofluorescence Imaging and Quantification

Fluorescent slides were imaged at 20× (Nanozoomer). QuPATH v0.4.4 (17) was used to quantify PCNA+ osteoblasts. Briefly, each half of the callus was manually contoured and Positive Cell Detection on the DAPI channel was used to identify the total number of nuclei. Single Measurement Classifiers were made for the PCNA and Osterix (Osx) signals individually, with thresholds based on maximum nuclear intensities for each channel. A Composite Measurement Classifier was created from the individual classifiers to identify PCNA-;Osx-, PCNA+;Osx-, PCNA-;Osx+, and PCNA+;Osx+ nuclei. Percent of proliferating osterix-expressing cells was calculated as the number of PCNA+;Osx+ cells divided by the total number of Osx+ nuclei. Similarly for Osteocalcin (Ocn)-stained sections, Positive Cell Detection on the DAPI channel was used to identify the total number of nuclei and the cell body was identified using cell expansion of 3 μm for 3-day and 4 μm for 5- and 10-day post-fracture samples. Osteocalcin thresholding was based on maximum cell intensity. Percent proliferating Ocn-expressing cells was calculated as the number of PCNA+;Ocn+ cells divided by the total number of Ocn+ cells. The two halves of each callus were averaged together to determine the rate of osteoblast proliferation throughout the entire callus (n=3 mice/timepoint).

#### Statistics

Experimenters were blinded to animal genotype and treatment during data collection and analysis. Analyses were carried out in GraphPad Prism 9.0. Data were tested for normality prior to analysis. Data were compared with Chi-Squared tests, unpaired two-tailed t-tests, one-way ANOVA with Holm-Sidak post-hoc correction, two-way ANOVA with Holm-Sidak post-hoc correction, or non-parametric Kruskal-Wallis multiple comparison tests, as indicated in figure legends. ANOVA tables are provided as a supplemental excel file for main variable effects. A p-value of <0.05 was used as a threshold for statistical significance. Graphs show means with error bars indicating SD.

---

**Figure S1.**
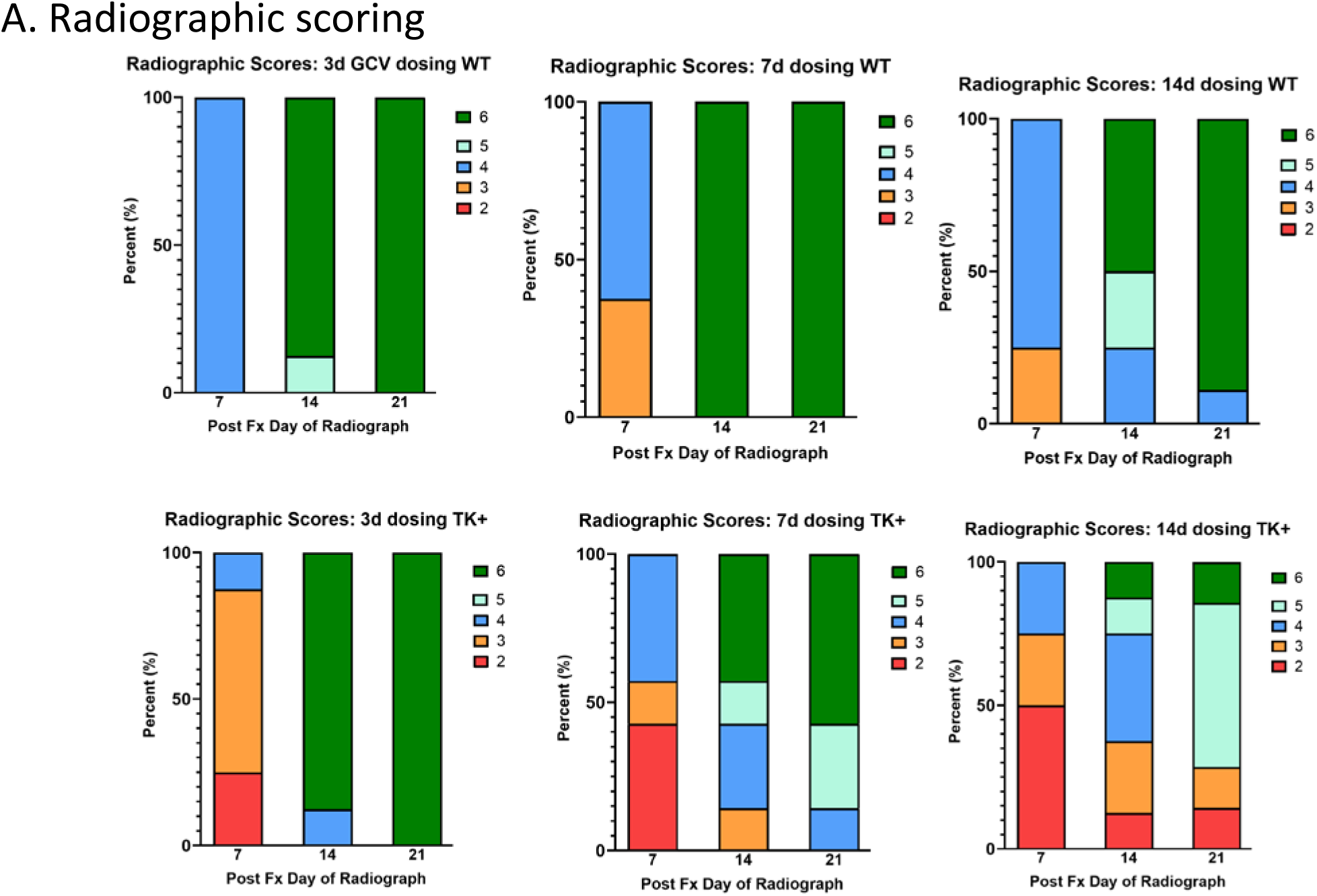
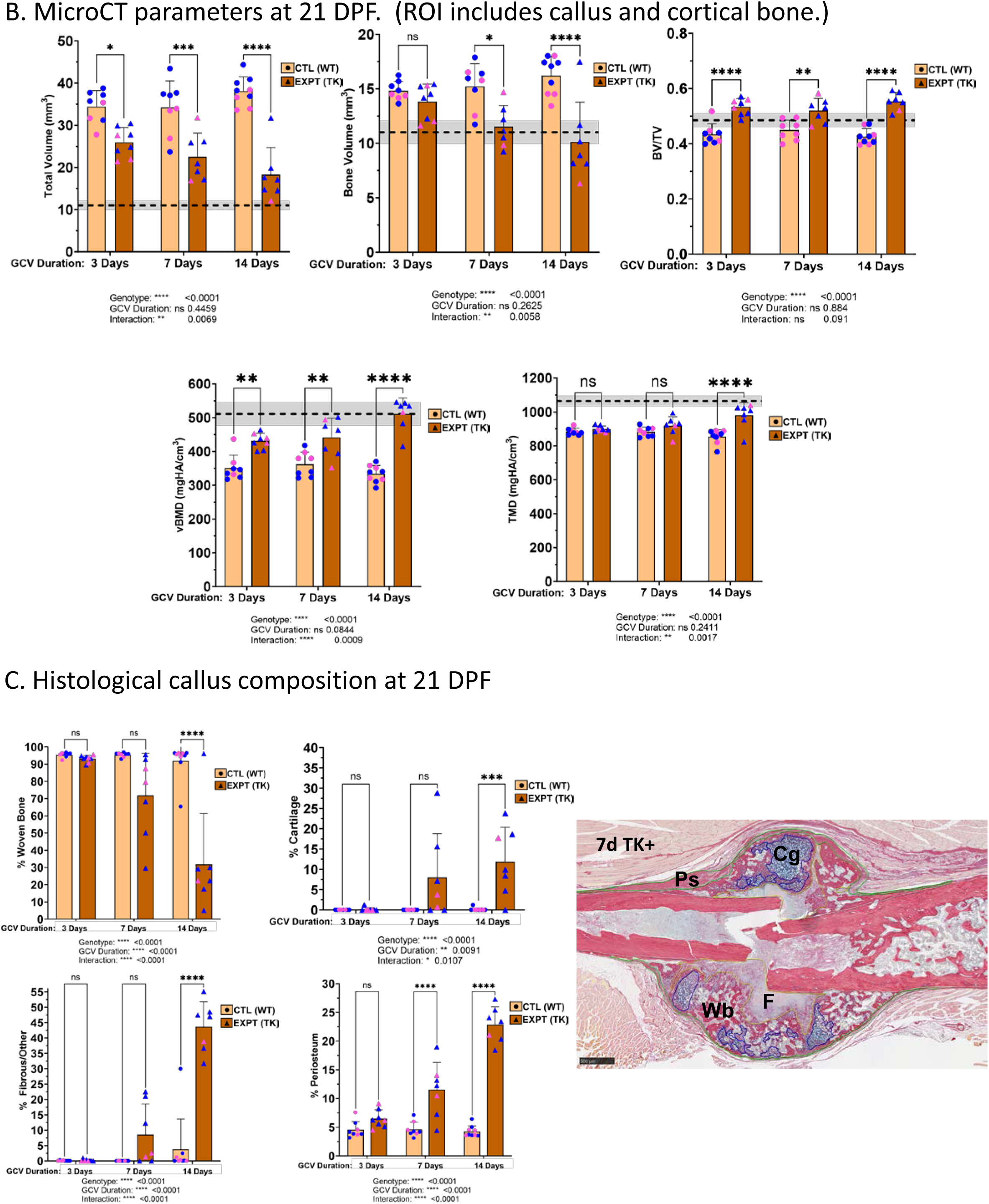
**(A)** Radiographic scoring by dosing duration at 7, 14 and 21 days post-fracture (DPF) (n=7-9). (Score 2=neither cortex bridged; 6=both cortices bridged). **(B)** microCT parameters from the callus region at 21 DPF (includes callus + cortical bone). (Dashed lines indicate avg. value of contralateral intact femurs; shading denotes +/- SD). **(C)** Histological callus composition at 21 DPF was determined from sections stained by picrosirius red and alcian blue (PSRAB). Image is representative section from Col1-TK mouse treated for 7 days with GCV. (Cg: cartilage, Wb: woven bone, F: fibrous tissue, Ps: periosteum.) (Graphs B, C depict mean ± SD; statistical differences determined by Fisher’s Exact Test)

**Supplemental Figure S2:**
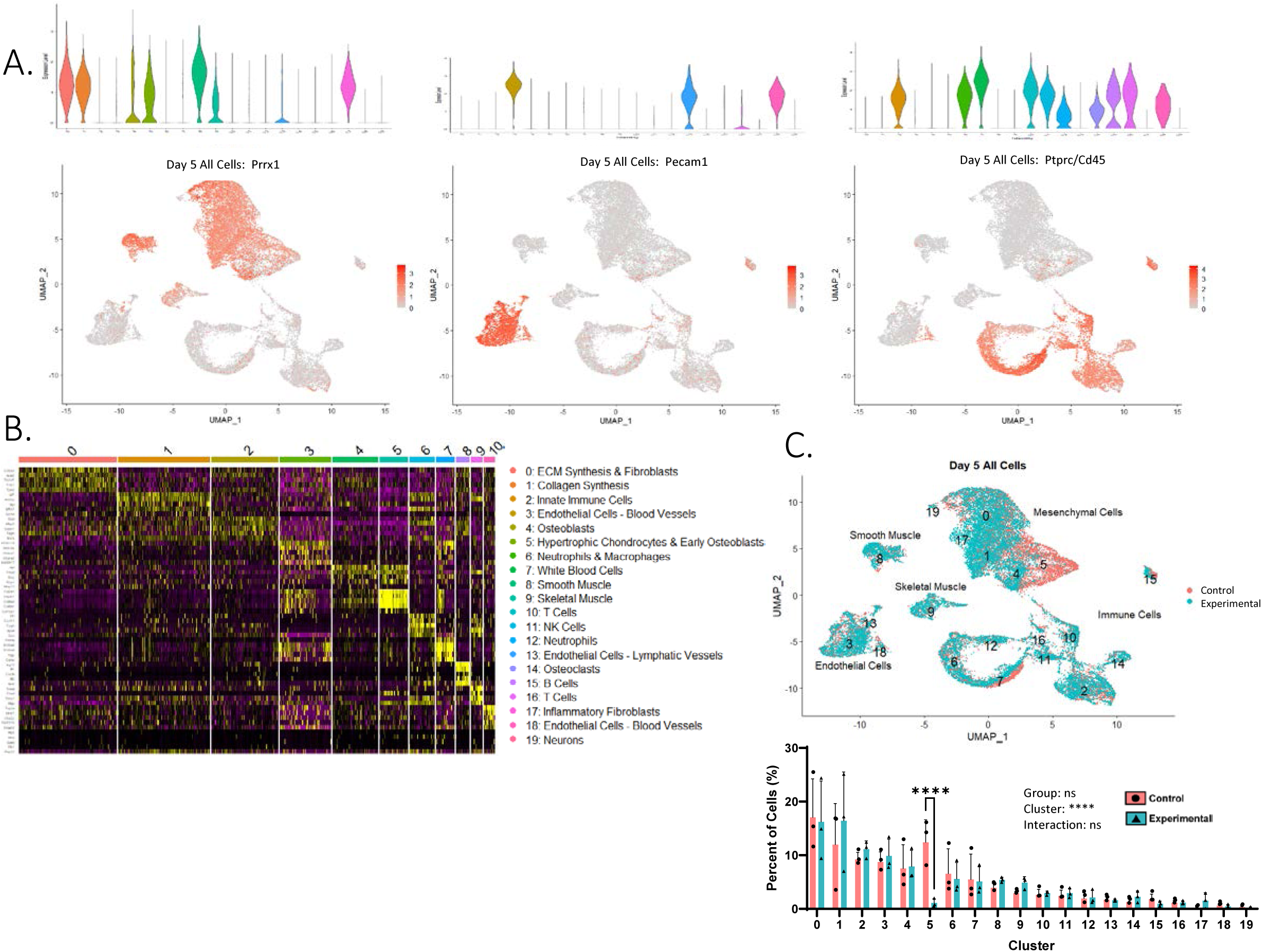
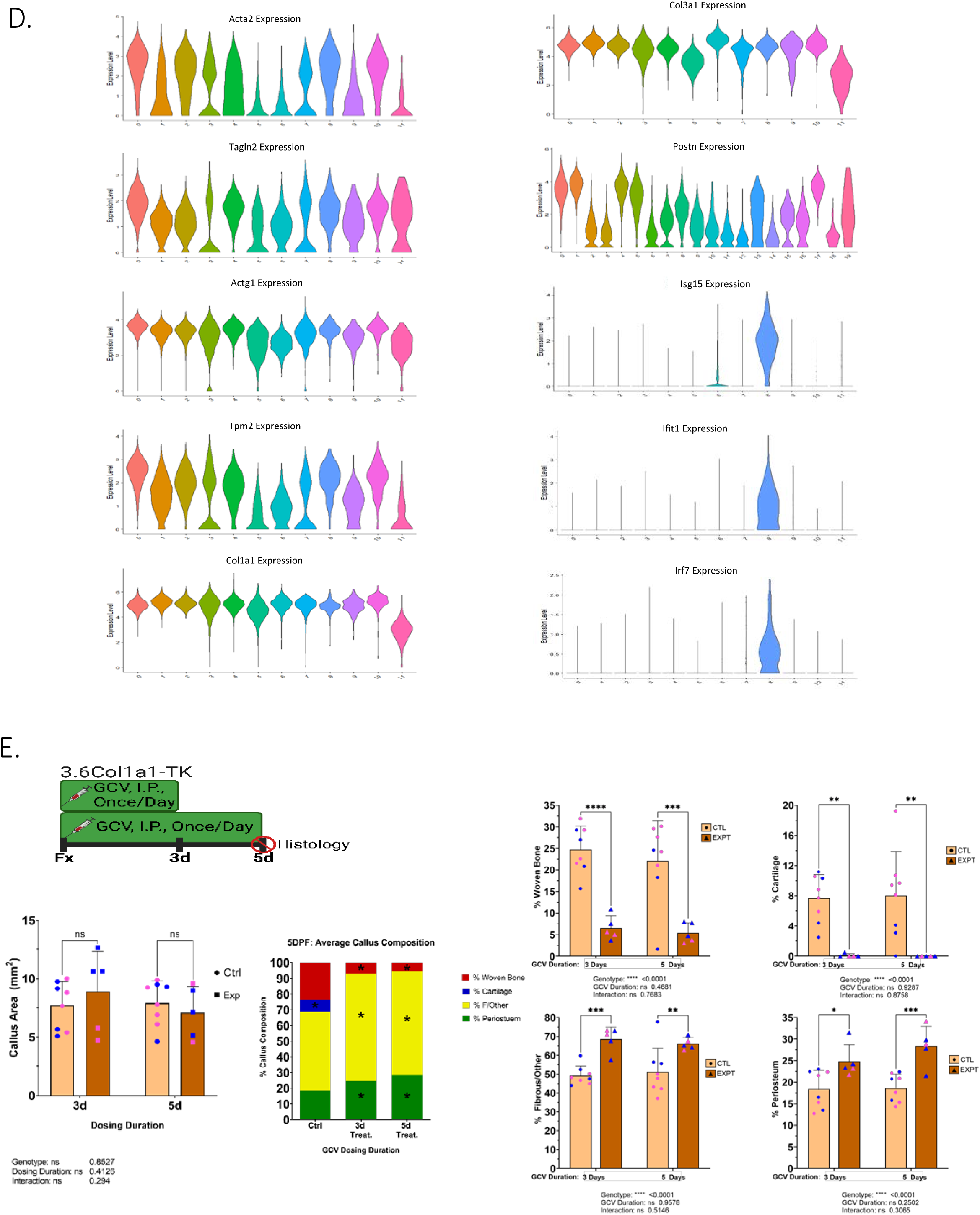
Ablating proliferating 3.6Col1a1-lineage cells during the first 5 days of healing alters fracture callus mesenchymal cell and tissue composition. **(A)** Expression of *Prrx1* (mesenchymal cells), *Pecam1* (endothelial cells), and *Ptprc*/CD45 (immune cells) in all cells 5 days post-fracture (DPF). **(B)** Top 5 DEGs for each cluster of all cells from 5 DPF. **(C)** UMAP colored by group (Control, Exptl). Percent of cells in each cluster by group was quantified (number of cells in cluster/total number cells from that callus) for each replicate callus and plotted (n=3). **(D)** Expression of myofibroblast genes (*Acta2*, *Tagln2*, *Actg1*, and *Tpm2*), nonspecific fibroblast genes (*Col1a1*,*Col3a1*, and *Postn*), and innate immune genes (*Isg15*, *Ifit1*, *Irf7*) used to annotate mesenchymal cell clusters. **(E)** Histological analysis of callus composition (n=5-8). (Graphs depict mean±SD; statistical differences were determined by two-Way ANOVA with Holm-Sidak post-hoc test.)

**Supplemental Figure S3:**
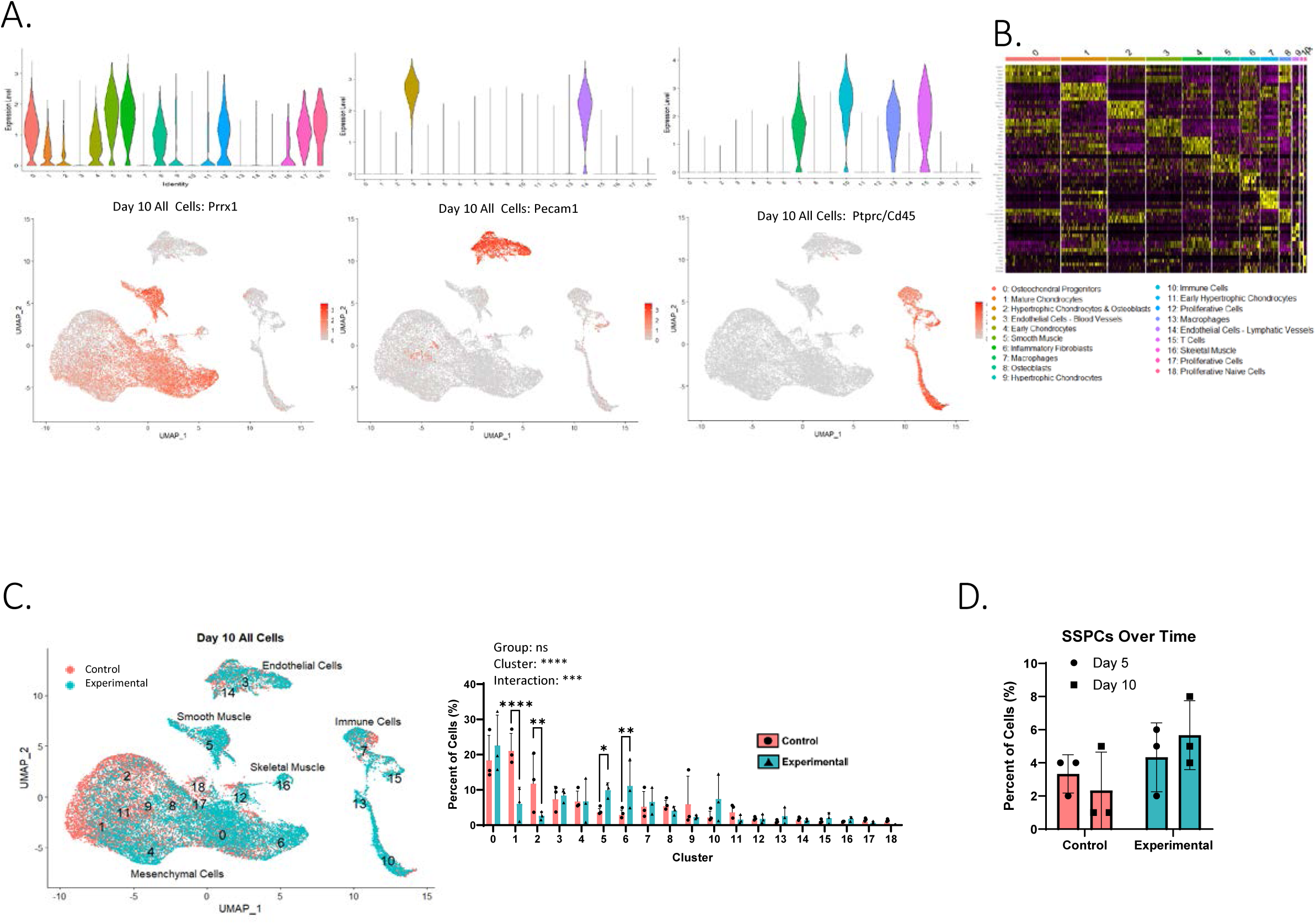
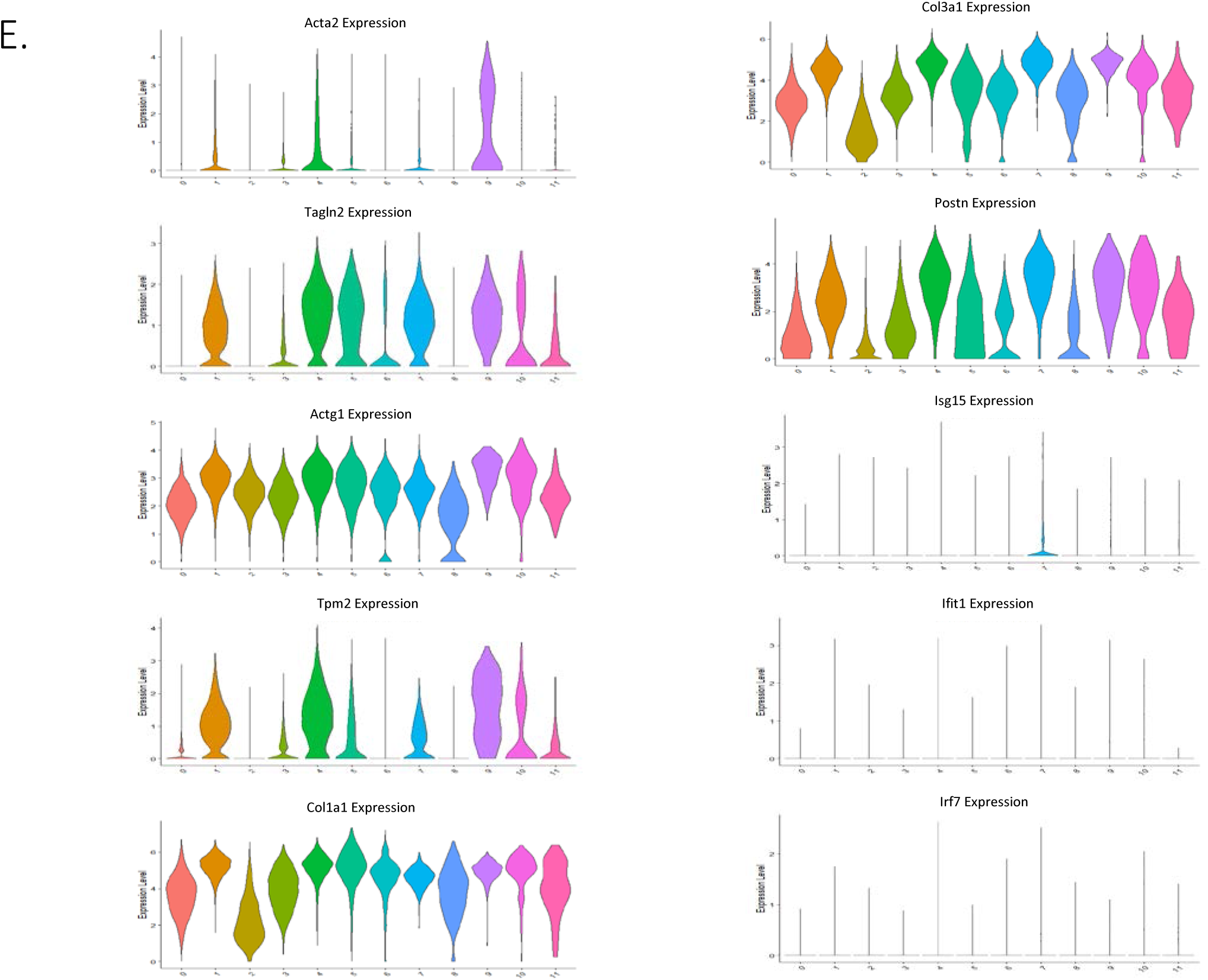
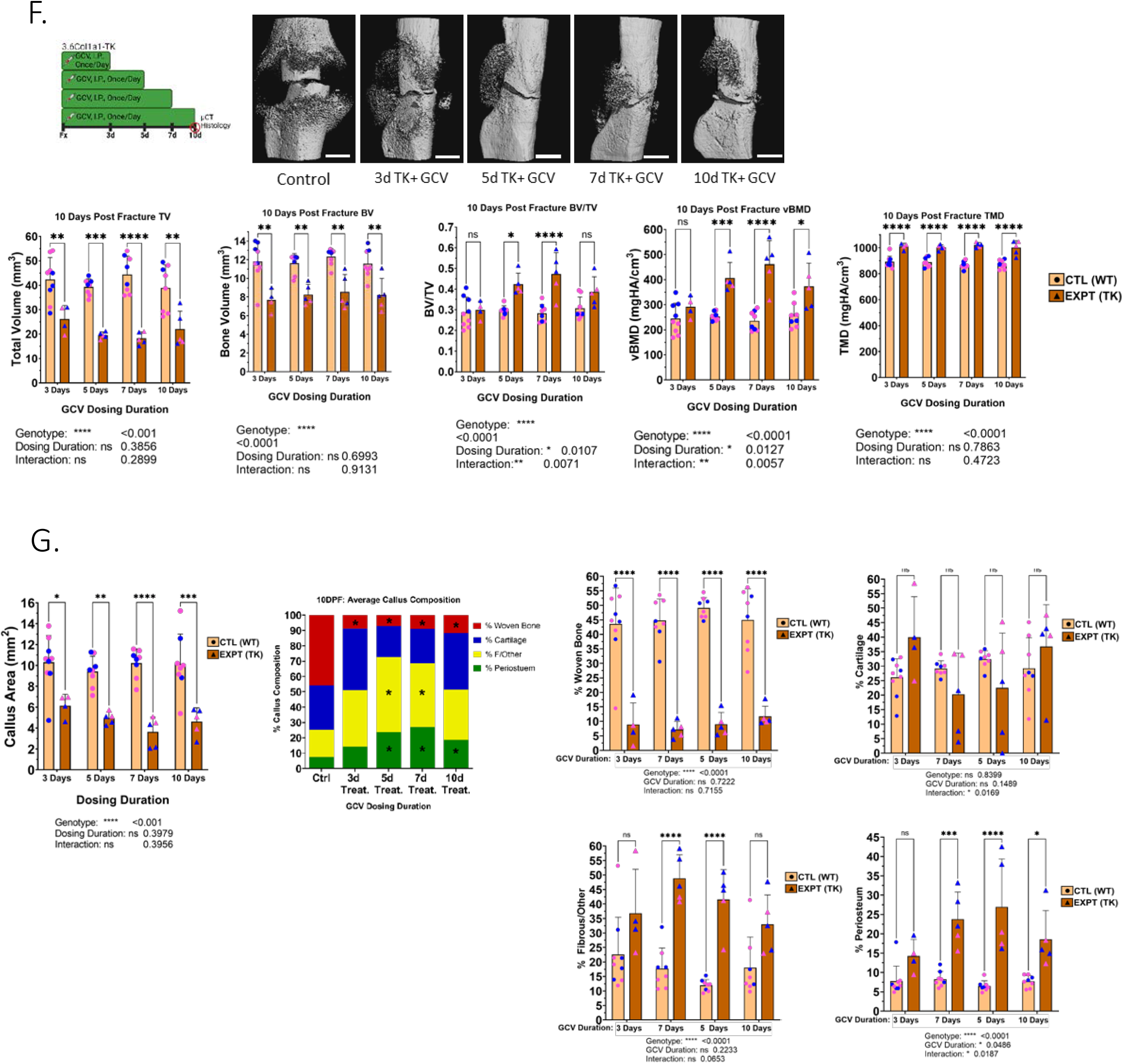
Ablating proliferating 3.6Col1a1-expressing cells during the first 10 days of healing significantly alters fracture callus mesenchymal cell and tissue composition. **(A)** Overlay of *Prrx1* (mesenchymal cells), *Pecam1* (endothelial cells), or *Ptprc*/CD45 (immune cells) of all cells 10 days post-fracture. **(B)** Top 5 DEGs for each cluster of all cells from 10 DPF. **(C)** UMAP colored by group (Control, Exptl). The percent of cells in each cluster by group was quantified (number of cells in cluster/total number of cells from that callus) for each replicate callus and plotted (n=3). **(D)** Number of Skeletal Stem and Progenitor Cells (SSPCs, defined as cells expressing *Acta2*, *Ly6a*, *Itgav*, *Thy1*, and *Ctsk*) normalized to the total number of cells from that replicate callus. **(E)** Expression of myofibroblast genes (*Acta2*, *Tagln2*, *Actg1*, and *Tpm2*), nonspecific fibroblast genes (*Col1a1*, *Col3a1*, and *Postn*), and innate immune genes (*Isg15*, *Ifit1*, *Irf7*) used to annotate mesenchymal cell clusters. **(F)** MicroCT analysis of fracture callus. **(G)** Histological analysis of callus composition. (Graphs depict mean±SD; statistical differences were determined by a two-Way ANOVA with Holm-Sidak Post-Hoc test.)

**Supplemental Figure S4:**
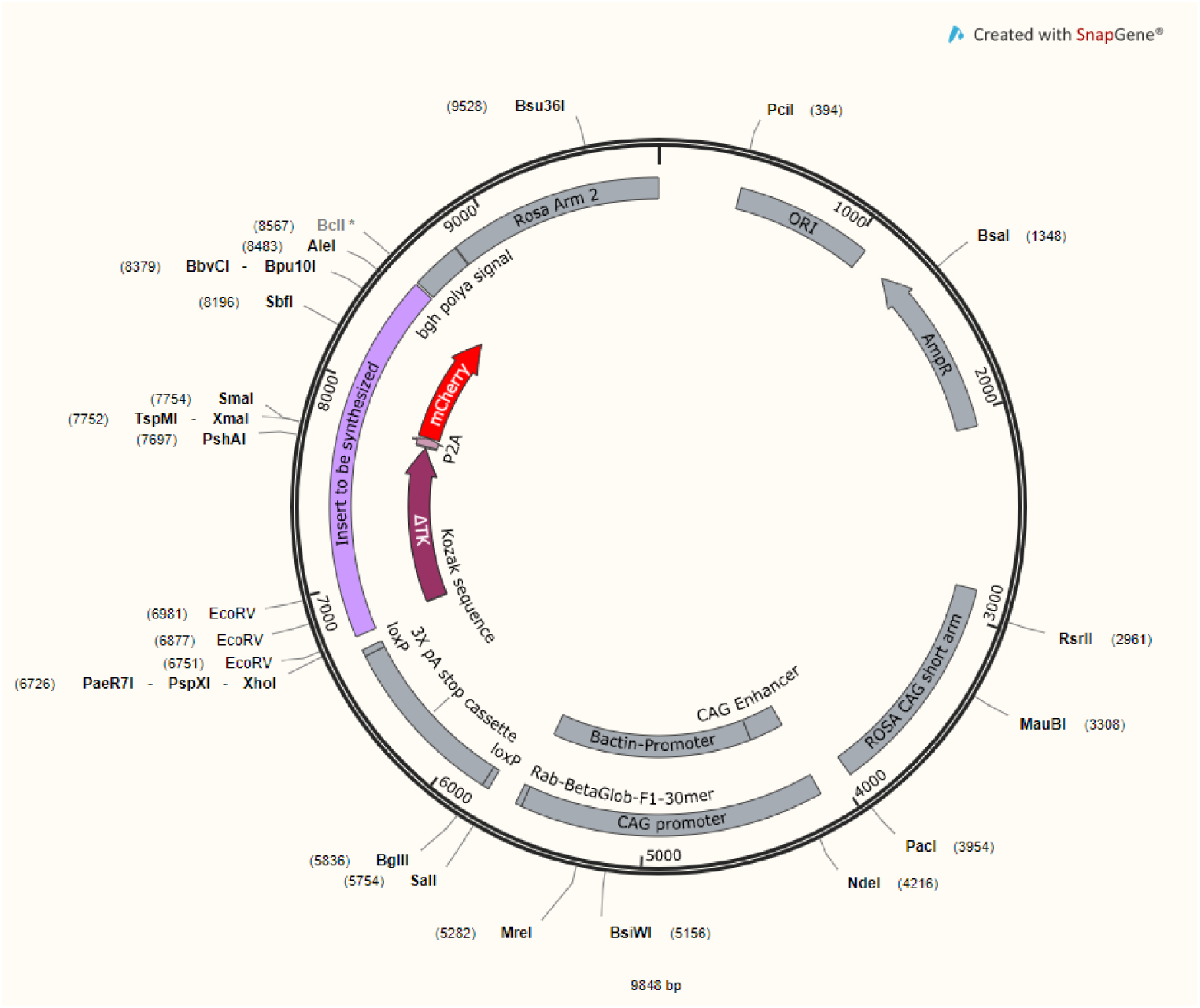
Targeting plasmid used to create ROSA-TK founders. Targeting plasmid that contains a CAG promoter, floxed stop cassette, the HSV-deltaTK allele, a P2A self-cleaving site, and an mCherry allele.

**Supplemental Figure S5:**
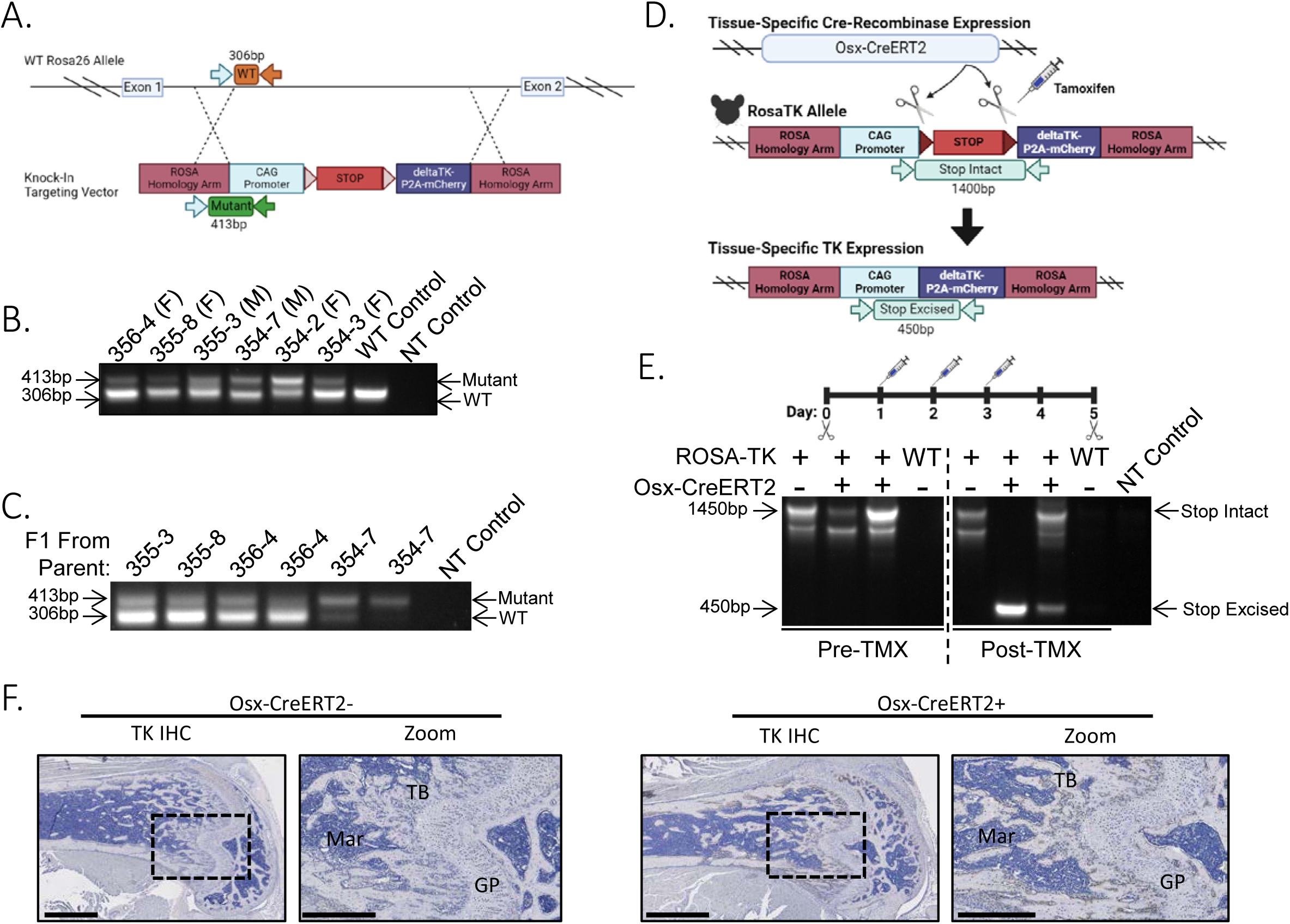
Generation of a novel ROSA-TK mouse line to drive tissue-specific TK expression using Cre-recombinase. **(A)** Schema of plasmid used to recombine the targeting vector into the ROSA26 allele. Genotyping primers to confirm proper recombination utilized a forward primer targeting the ROSA26 homology arm and reverse primers targeting either the wild-type ROSA26 allele or the mutant CAG promoter. **(B)** Six heterozygous founders were identified using the above-described primers and were then bred with wild-type B6 mice. **(C)** Progeny from each of the six founders maintained the mutant allele. **(D)** Genotyping scheme to evaluate stop cassette excision in response to tamoxifen (TMX) activation of inducible Osx-CreERT2. Genotyping primers spanning either the intact stop cassette or the CAG promoter and TK gene delineate the intact versus excised stop cassette. **(E)** Tail snips from Osx-CreERT2;ROSA-TK mice prior to TMX treatment shows only intact stop cassettes. After three TMX treatments, a second tail snip from the same mice shows stop cassette excision only in mice that have both ROSA-TK and Osx-CreERT2 alleles. Mice lacking Osx-CreERT2 show retention of the stop cassette and mice without the ROSA-TK allele show no PCR amplification of either the intact or excised stop cassette products. **(F)** IHC for TK shows that TMX induces TK expression in osteoblasts lining the trabecular bone in the growth plate in 5-week-old Osx-CreERT2;ROSA-TK mice but not Osx-CreERT2-(neg); ROSA-TK mice. Scale bar = 1 mm in TK IHC, 0.5 mm in Zoom.

**Supplemental Figure S6:**
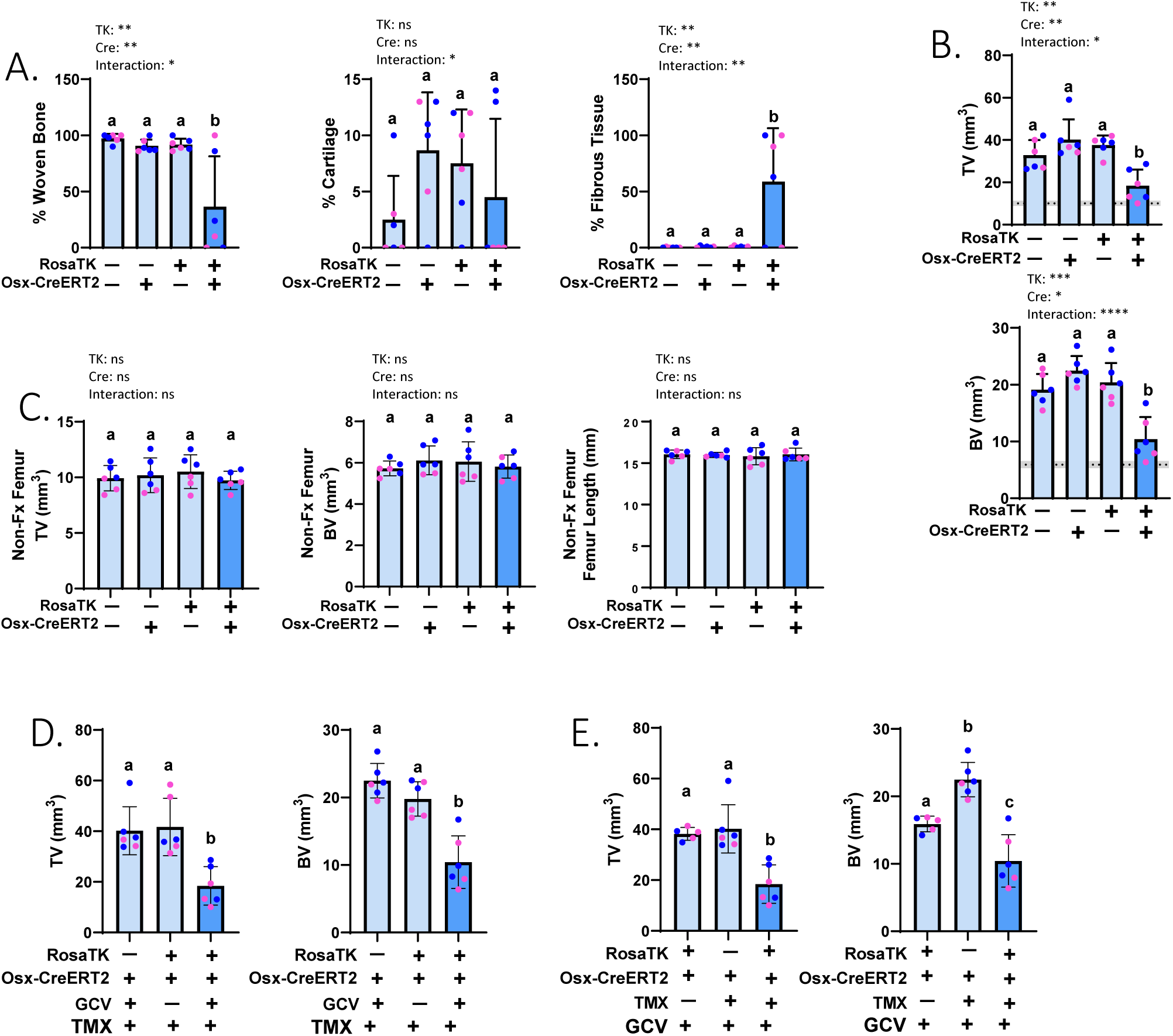
Ablation of proliferating Osterix-expressing cells significantly impairs fracture callus mineralization without affecting the contralateral non-fractured limb. **(A)** Histological analysis of callus. Ablation of proliferating Osterix-expressing cells for 14 days post-fracture significantly reduces the amount of woven bone and significantly increases the amount of fibrous tissue within the fracture callus, without affecting %cartilage. These data are summarized in the stacked bar graph in Figure 4D. **(B)** microCT analysis of the callus region (includes callus + cortical bone). Ablation of proliferating Osterix-expressing cells for 14 days post-fracture significantly reduces Total Volume (TV) and Bone Volume (BV) at the site of fracture callus. (Dashed lines indicate avg. value of contralateral intact femurs; shading denotes +/- SD). **(C)** microCT analysis of non-fractured bones. Treatment with GCV does not significantly change TV, BV, or femur length in the contralateral, non-fractured femurs. (All mice in A-C were dosed with GCV.) **(D)** Impairments in fracture callus size and bone volume are only seen in mice that are Cre+;TK+ and treated with GCV, even if all received TMX treatment. **(E)** Impairments in fracture callus size and bone volume are only seen in mice that are Cre+;TK+ and are treated with TMX, even if all received GCV treatment. (The third bar of graphs in panels D and E show identical data, also shown in the fourth bar of panel B.) (Graphs depict mean±SD and statistical differences were determined by (A-C) a Two-Way ANOVA with Holm-Sidak Post-Hoc test or (D & E) Kruskal-Wallis test.)

**Supplemental Figure S7:**
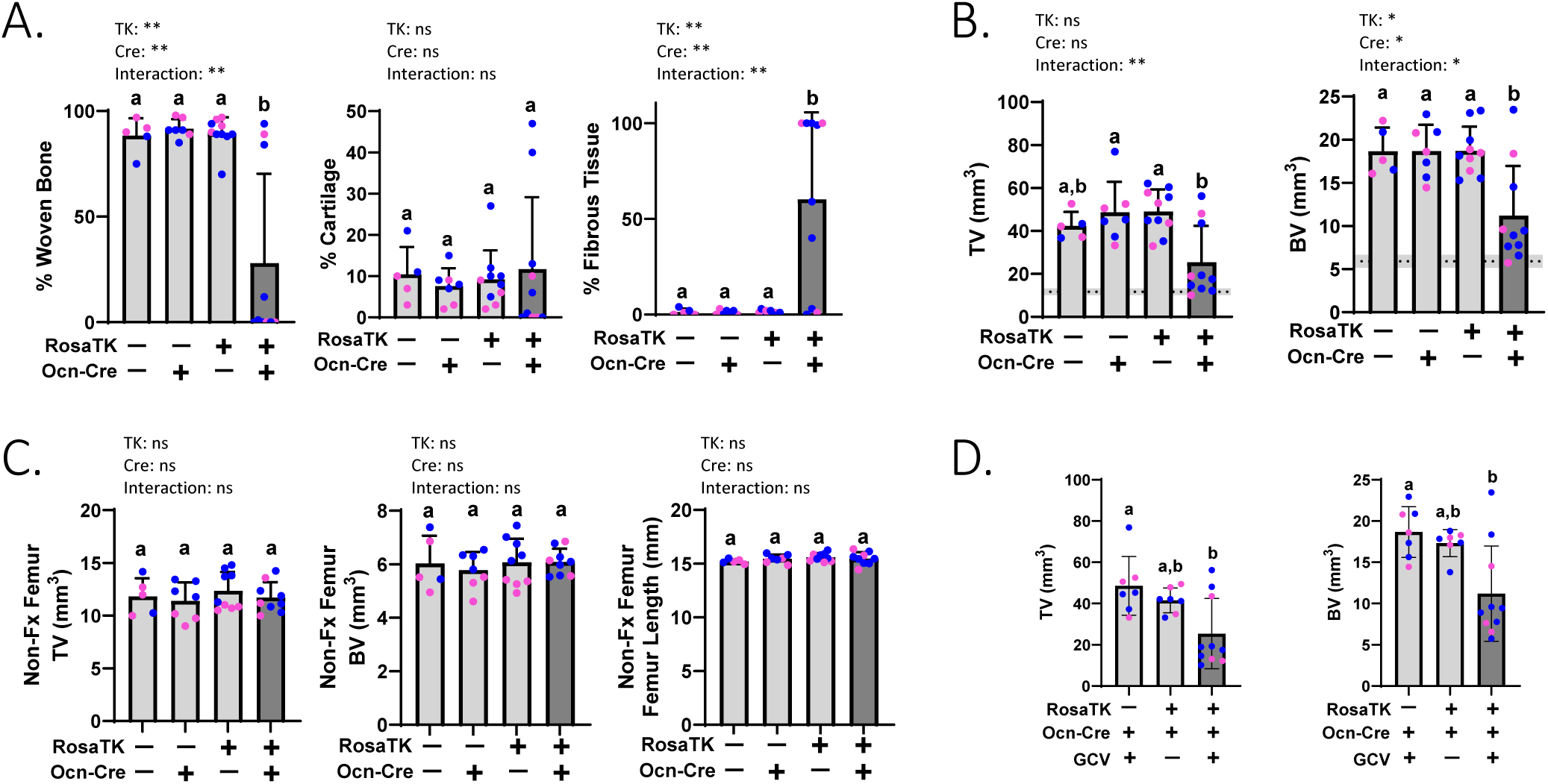
Ablation of proliferating Osteocalcin-lineage cells significantly impairs fracture callus mineralization without affecting the contralateral non-fractured limb. **(A)** Fracture callus composition was determined from histology. TK+/Ocn-Cre+ mice have reduced woven bone and increased fibrous tissue within the fracture callus without affecting cartilage composition. These data are summarized in the stacked bar graph in Figure 6D. **(B)** MicroCT analysis of callus region (includes callus + cortical bone). TK+/OcnCre+ mice have reduced Total Volume (TV) and Bone Volume (BV). (Dashed lines indicate avg. value of contralateral intact femurs; shading denotes +/- SD). **(C)** MicroCT of intact femurs. Genotype does not alter TV, BV, or Femur Length in the contralateral, non-fractured femurs from the same cohort of mice. (All mice in A-C were dosed with GCV.) **(D)** Impairments in fracture callus morphology are only seen in mice that are Cre+;TK+ and treated with GCV. (Graphs depict mean±SD and statistical differences were determined by a two-Way ANOVA with Holm-Sidak Post-Hoc test.)

**Supplemental Figure S8:**
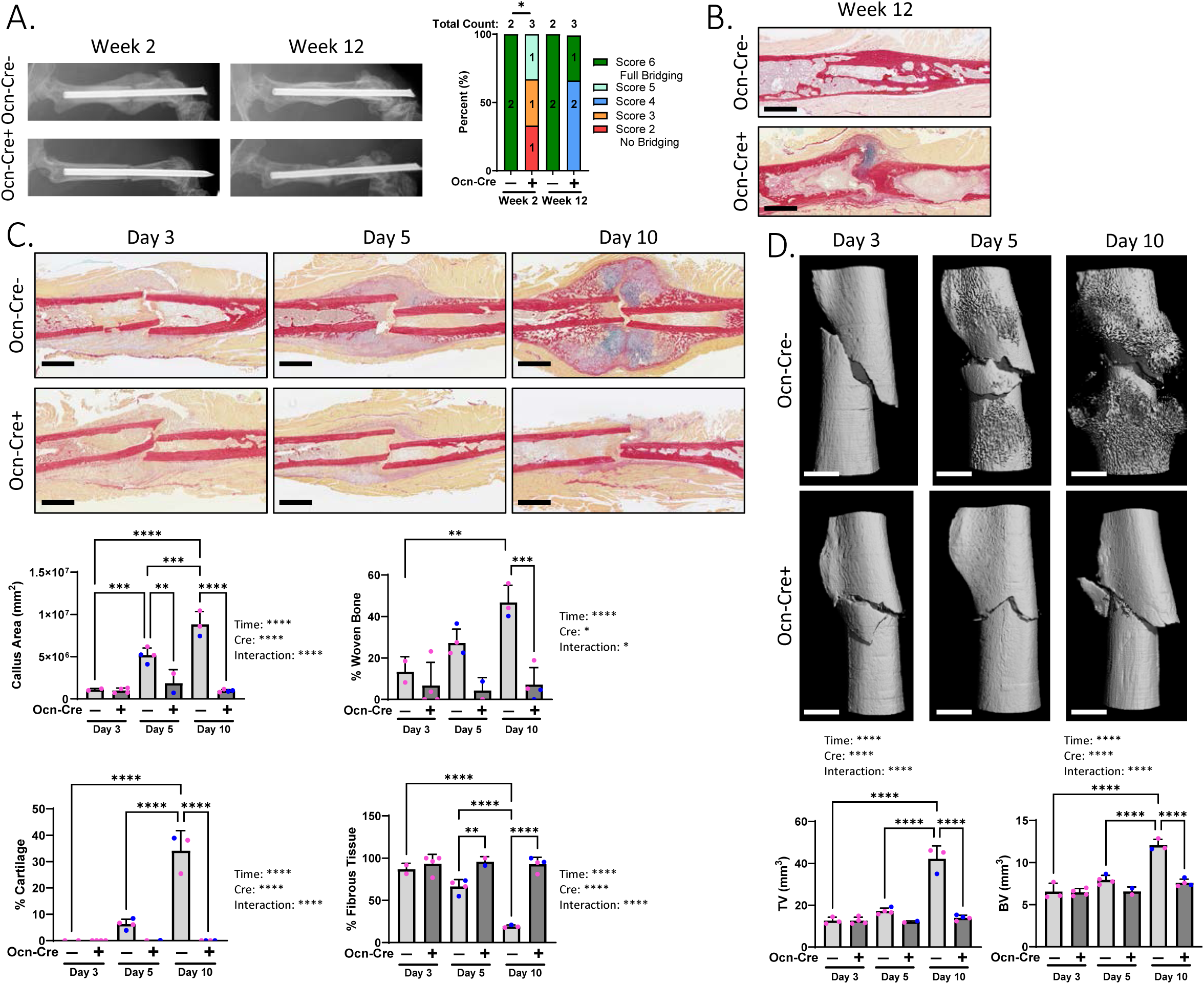
Ablation of proliferating Osteocalcin-lineage cells significantly impairs fracture callus healing in long- and short-term. **(A)** Ocn-Cre;ROSA-TK mice were treated with GCV for 2 weeks post-fracture (as depicted in Fig 6A), then GCV was withdrawn for the following 10 weeks. X-ray scoring shows that Ocn-Cre-mice fully bridge and remodel by 12 weeks post-fracture, whereas Ocn-Cre+ mice have significantly impaired bridging at 2 weeks post-fracture that does not fully recover by 12 weeks. **(B)** PSRAB staining of representative samples shows that Ocn-Cre-mice 12 weeks post-fracture have bony healing at the fracture site, whereas Ocn-Cre+ mice still have cartilaginous and fibrotic calluses 12 weeks post-fracture. **(C)** Ocn-Cre;ROSA-TK mice were treated with GCV for 3-, 5-, or 10-days post-fracture and euthanized on the last day of treatment indicated. PSRAB staining and analysis shows early changes in callus composition. **(D)** microCT analysis of Ocn-Cre;RosaTK mice treated with GCV for 3-, 5-, or 10-days post-fracture and euthanized on the last day of treatment indicated shows that control Cre-mice increase callus mineralization over time, whereas Cre+ experimental mice do not create a mineralized callus. (Graphs depict mean±SD; statistical differences were determined by (A) Chi-Square test, or (C-D) two-Way ANOVA with Holm-Sidak Post-Hoc test.)

**Supplemental Figure S9:**
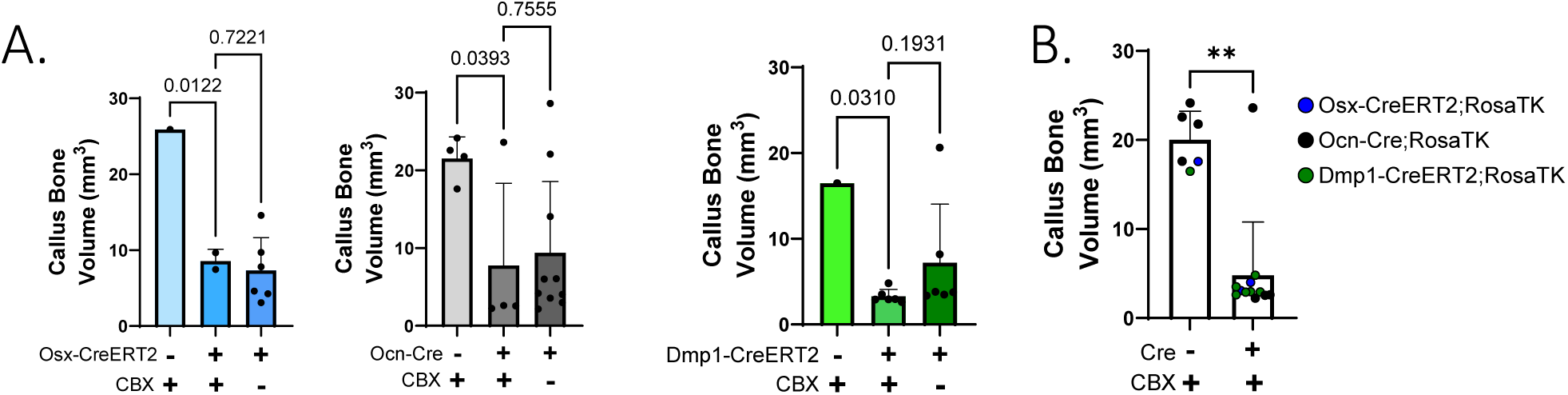
Treatment with gap junction inhibitor does not rescue impaired fracture healing in Osx-CreERT2, Ocn-Cre, or Dmp1-CreERT2;ROSA-TK mice 2 weeks post-fracture. **(A)** Cre;ROSA-TK mice were treated with GCV and also treated with carbenoxolone (CBX) to block gap junction intercellular communication. In three lines of mice receiving CBX, callus bone volume measured by microCT was significantly less in Cre+ experimental vs. Cre-control. Callus bone volume in Cre+/CBX+ mice was not different from experimental Cre+ mice that did not receive CBX (the latter data also shown in Figs. 4E, 6E, and 8E). **(B)** When data from each Cre line were aggregated to overcome low sample size, Cre+ mice had impaired callus mineralization compared to Cre-mice, both treated with CBX. (Graphs depict mean±SD; statistical differences were determined by One-Way ANOVA with Fisher’s LSD Post-Hoc (A) or Mann-Whitney U test (B).)

**Supplemental Figure S10:**
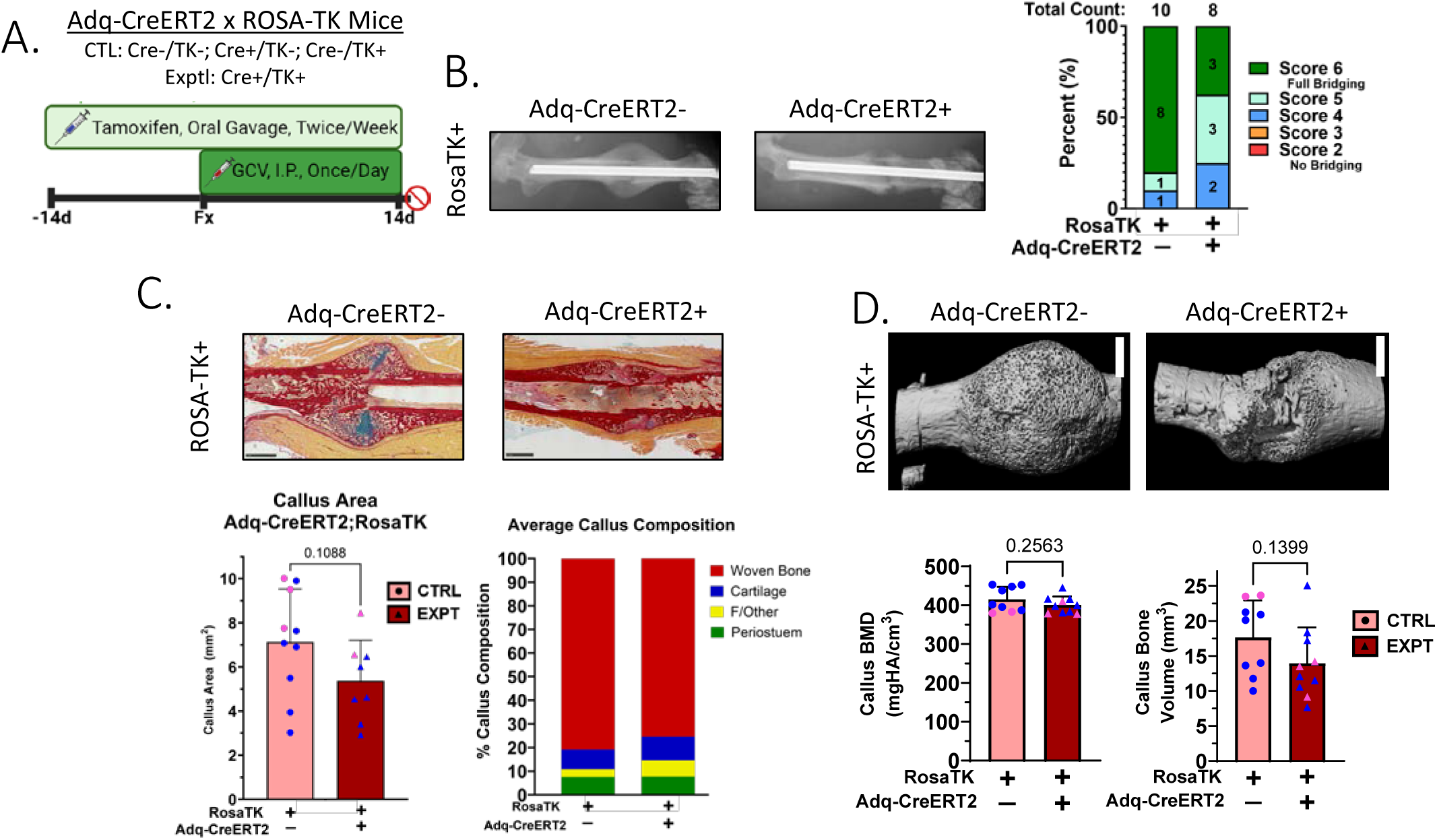
Ablation of proliferating adiponectin expressing cells modestly affects fracture healing. **(A)** Mice were treated with TMX starting at 10-wks age, followed by femur fracture at 12-wks, followed by 2 weeks GCV treatment. Healing was assessed 14 days after fracture. Control mice were Cre-;TK+ while experimental mice were Cre+;TK+. **(B)** Radiographs showed less mineralized callus in Cre+ mice, resulting in modestly poorer callus bridging scores than in Cre-controls. **(C)** Histological analysis shows a marginally smaller callus area in Cre+ mice, but normal percent cartilage and woven bone compared to Cre-controls. **(D)** Fractured bones were assessed by microCT; there were no significant differences in callus bone volume or BMD between Cre+ mice and control mice. (Statistical differences were determined by Chi-Square test (B) or by two-tailed t-test (C,D). Scale bars: PSRAB=1 mm, TK IHC=0.5 mm, microCT=1 mm.)

